# Large pilin subunits provide distinct structural and mechanical properties for the *Myxococcus xanthus* type IV pilus

**DOI:** 10.1101/2023.07.22.550172

**Authors:** Anke Treuner-Lange, Weili Zheng, Albertus Viljoen, Steffi Lindow, Marco Herfurth, Yves F. Dufrêne, Lotte Søgaard-Andersen, Edward H. Egelman

**Affiliations:** Max Planck Institute for Terrestrial Microbiology, 35043 Marburg, Germany; Department of Biochemistry and Molecular Genetics, University of Virginia School of Medicine, Charlottesville, VA 22903, USA; Louvain Institute of Biomolecular Science and Technology, UCLouvain, B-1348 Louvain-la-Neuve, Belgium

## Abstract

Type IV pili (T4P) are ubiquitous bacterial cell surface filaments important for surface motility, adhesion to biotic and abiotic surfaces, DNA uptake, biofilm formation, and virulence. T4P are built from thousands of copies of the major pilin subunit and tipped by a complex composed of minor pilins and in some systems also the PilY1 adhesin. While the major pilins of structurally characterized T4P have lengths of up to 161 residues, the major pilin PilA of *Myxococcus xanthus* is unusually large with 208 residues. All major pilins have a highly conserved N-terminal domain and a highly variable C-terminal domain, and the additional residues in the *M. xanthus* PilA are due to a larger C-terminal domain. We solved the structure of the *M. xanthus* T4P (T4P^Mx^) at a resolution of 3.0 Å using cryo-electron microscopy (cryo-EM). The T4P^Mx^ follows the structural blueprint observed in other T4P with the pilus core comprised of the extensively interacting N-terminal α1-helices while the globular domains decorate the T4P surface. The atomic model of PilA built into this map shows that the large C-terminal domain has much more extensive intersubunit contacts than major pilins in other T4P. As expected from these greater contacts, the bending and axial stiffness of the T4P^Mx^ is significantly higher than that of other T4P and supports T4P-dependent motility on surfaces of different stiffnesses. Notably, T4P^Mx^ variants with interrupted intersubunit interfaces had decreased bending stiffness and strongly reduced motility on all surfaces. These observations support an evolutionary scenario whereby the large major pilin enables the formation of a rigid T4P that expands the environmental conditions in which the T4P system functions.

## Introduction

Bacterial motility is important for virulence, colonization of various habitats, biofilm formation, interactions with host cells, and fitness by directing cells toward nutrients and away from toxins and predators^1^. Accordingly, bacteria can move in many different environments and their motility devices are adapted to these varying conditions^1, 2^. Generally, bacteria move using highly conserved nanomachines that energize either the rotation of flagella to enable swimming in liquids and swarming on semisolid surfaces or the extension/retraction of type IV pili (T4P) to enable translocation on solid surfaces^1, 2^. Some bacteria also move on surfaces by gliding, but the involved nanomachines are more diverse, each with a narrow taxonomic distribution^1, 2^. Here, we focus on T4P, which are not only important for motility but also for adhesion to host cells and abiotic surfaces, natural transformation with horizontal gene transfer, biofilm formation, virulence, predation, and surface sensing^3, 4^.

The versatility of T4P depends on their ability to undergo cycles of extension with adhesion to a surface, and retractions that generate a force sufficient to pull a cell forward^5–8^. These cycles are driven by the T4P machine (T4PM), which is composed of ∼15 conserved proteins forming a large macromolecular complex that spans from the outer membrane across the periplasm and inner membrane (IM) to the cytoplasm^9–11^. T4P extension and retraction are powered by the ATPases PilB and PilT, respectively that bind to the cytoplasmic base of the T4PM in a mutually exclusive manner^3, 10, 12^. All ∼15 proteins are essential for T4P extension except for PilT, which is only necessary for retraction^3^. The T4P are flexible, thin, up to several microns in length, and composed of thousands of copies of the major pilin subunit as well as a tip complex comprising minor pilins and sometimes also the PilY1 adhesin^11, 13–16^. During T4P extension, major pilin subunits are extracted from a reservoir in the IM and inserted at the base of the growing pilus; during T4P retractions, this process is reversed, and the major pilin subunits are removed from the base of the T4P and reinserted into the IM^3^.

Major pilins are synthesized as prepilins with an N-terminal type III signal peptide (T3SP), which is cleaved off by the PilD prepilin peptidase between the Gly and Phe residues in the consensus GFxxxE motif to generate the mature major pilin (hereafter simply referred to as the major pilin)^17^. Sequence and structural analyses of major pilins in isolation and structural studies of intact T4P filaments have shown that major pilins share the same overall structure with a semi-conserved N-terminal α-helix (α1) and a highly variable C-terminal, largely β-stranded, globular domain^4, 13^. In Gram-positive bacteria, the C-terminal domain can be all α-helical^18^, while in *Geobacter sulfurreducens* it has now been shown that the two domains are encoded by two different genes^19^, resulting in a pilin subunit containing two polypeptide chains. Proteins homologous to major pilins are also the building blocks of the endopilus (previously called pseudopilus) of the type II secretion system as well as archaeal T4P and flagella^4^. Since in archaea both T4P pilins and flagellins have homology with only the N-terminal bacterial pilin domain, the suggestion has been made that all pilins have arisen from a gene fusion of ancestral genes encoding the N-and C-terminal domains separately^20^. The α1-helix can be divided into the mainly hydrophobic highly-conserved N-terminal part (α1-N), which is essential for anchoring the pilin in the IM before its incorporation into the pilus, and the less conserved amphipathic C-terminal part (α1-C) that connects to and packs against the globular C-terminal domain^4, 13^.

The structures of eight bacterial T4P filaments, including an endopilus of a type II secretion system, have been solved to a resolution of 3.2-8.0 Å, from *Neisseria gonorrhoeae* (PDB 5VXX)^21^, *N. meningitidis* (PDB 5KUA)^22^*, Pseudomonas aeruginosa* (PDB 5VXY)^21^*, Escherichia coli* (PDB 6GV9)^23^, *G. sulfurreducens* (PDB 6VK9 and 7TGG)^19, 24^, *Klebsiella oxytoca* (PDB 5WDA; endopilus)^25^ and two different ones from *Thermus thermophilus* (PDB 6XXD and 6XXE)^26^. These structures revealed, not surprisingly, that all T4P filaments share the same overall architecture. Specifically, the major pilins are helically arranged and tightly packed, giving rise to pili with widths of ∼60-75 Å, a rise of ∼9-11 Å per subunit, and ∼4 subunits per turn, with ∼1000 subunits per micron length of the T4P. The pilus core comprises the extensively interacting α1-helices while the variable globular domains decorate the T4P surface. In this conserved structural blueprint, the α1-helices establish the backbone of the T4P, while the divergent globular domains determine the shape, surface charge, and functional properties of T4P^21–23, 25–27^. While the C-terminal globular domain remains largely unchanged upon the incorporation of a major pilin into the pilus, the α1-helix undergoes a partial loss of α-helical structure of variable length around the highly conserved Pro22 residue^19, 21, 22, 26^. It was suggested that the melting of this segment is essential for the tight packing of the major pilins^21^. The extensive interactions between major pilins make T4P highly robust, and in the case of *N. gonorrhoeae* and *M. xanthus,* T4P were shown to withstand pulling forces of 110 to 150pN, respectively, during retractions^7, 8^. In addition, T4P of *N. gonorrhoeae, N. meningitidis* and *P. aeruginosa* have been shown to be highly extensible, undergoing force-induced conformational changes to elongate in response to pulling forces^28–31^. Moreover, in *N. gonorrhoeae,* these conformational changes were shown to be reversible^29^. Whether this resilience is a conserved feature of T4P is not known. It was proposed^21^ that further melting of the N-terminal α-helix is responsible for the extensibility of these filaments, and the restoring force after extension would be provided by the refolding of the α-helix.

Major pilins are not only diverse in sequence but also in size^4^. Among the solved T4P structures, the major pilins vary in size from 111-161 residues^21–23, 26^, while the two polypeptide chains forming the *G. sulfurreducens* pilin have a combined size of 165 residues^19, 24^. *M. xanthus* is a model system for understanding the architecture and mechanism of the T4PM^10, 11^. Of note, the major pilin PilA of *M. xanthus* contains 208 residues^32^ and is, thus, significantly larger than those of solved T4P structures. Moreover, the *M. xanthus* T4P (henceforth T4P^Mx^) is highly robust and can withstand a pulling force of 150pN during retractions^7^. *M. xanthus* is a predatory soil bacterium and belongs to the myxobacteria, prolific secondary metabolite producers^33^. *M. xanthus* has a biphasic nutrient-regulated lifestyle in which cells organize to form spreading, predatory colonies in the presence of nutrients and spore-filled fruiting bodies in the absence of nutrients^34, 35^. In both phases of the lifestyle, motility has a key function. *M. xanthus* has two motility systems for translocation across surfaces, one for gliding and one that depends on T4P^34, 35^. These two motility systems enable *M. xanthus* cells to translocate on highly diverse surfaces^36^.

To understand the properties conferred by large major pilins to T4P filaments, we determined the structure of T4P^Mx^ using cryo-EM to a resolution of 3.0 Å, which allowed us to build *de novo* an atomic model of the entire major pilin, and analyzed its biophysical properties *in vitro.* This structure revealed a T4P that differed from all existing T4P structures since it had much more extensive contacts between the globular domains. Consistent with such a structure, the stiffness of T4P^Mx^ is significantly higher than that of *N. gonorrhoeae* and *P. aeruginosa* T4P. Structure-guided mutagenesis of PilA showed that disruption of subunit interfaces caused a reduction in the axial stiffness of the T4P, and these variant T4P were less efficient in supporting motility on surfaces of different stiffness.

## Results

### Major pilins vary significantly in size

To systematically assess the size of major pilins, we first extracted all sequences of the K02650 group (type IV pilus assembly protein PilA) from the KEGG Orthology (KO) database^37^. After filtering out sequences with >90% sequence identity, sequences lacking a T3SP, and/or sequences lacking a classified taxonomy, we obtained a set of 1,955 prepilins of T4P. After removal of the T3SP, the major pilins vary in length from 42 to 297 aa, with a mean of 141±25 aa, in good agreement with a previous estimate based on fewer sequences^13^ (Fig. 1A, Table S1). Because the largest structurally characterized major pilin has a size of 161 aa (and 165 aa for the heterodimeric *G. sulfurreducens* major pilin), we arbitrarily defined large major pilins as proteins with a size ≥166 aa. Among our set of 1,955 sequences, 226 proteins, representing 12%, fulfilled this criterion. These proteins are widespread and present in 13 of the 21 phyla with major pilins, and largely group according to phylogeny (Fig. 1A; Fig. S1; Table S1). However, their distribution in phyla and classes is highly skewed, and at the phyla and class levels, they are overrepresented in Betaproteobacteria (20%), Cyanobacteria (23%), Myxococcota (93%) and Bdellovibrionota (100%) (Fig. 1A, Table S1). Moreover, while the length distribution of major pilins in Betaproteobacteria (42-279 aa) and Cyanobacteria (105-243 aa) is broad (Fig. 1A), it is more narrow in the predatory Myxococcota (153-217 aa) and Bdellovibrionota (170-204 aa) (Fig. 1A). Interestingly, in the Betaproteobacteria, the large major pilins are enriched explicitly in the order Burkholderiales (66 of 75) (Table S2), an ecologically diverse order that includes plant, animal and human pathogens^38^, and especially in the *B. cepacia* complex, which is associated with cystic fibrosis^39, 40^. Similarly, in the Firmicutes, the large mature major pilins are highly enriched in the order Eubacteriales (19 of 25), which includes several species of the gut microbiome (Table S2) in which T4P are ubiquitous^41, 42^.

**Figure 1.**
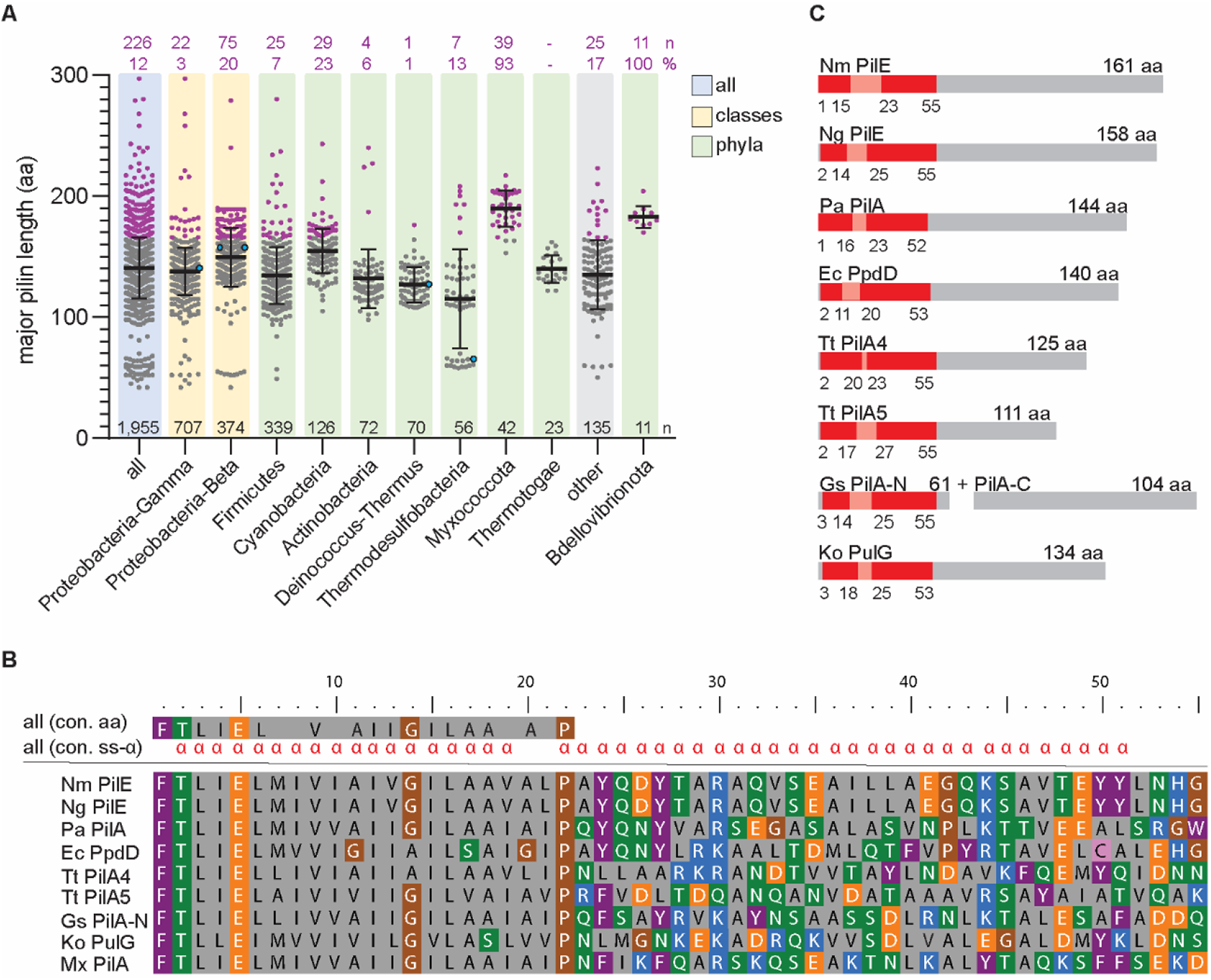
Length and phylogenetic distribution of major pilins. **A.** The length distribution of major pilins of the KEGG orthology group K02650 (type IV pilus assembly protein PilA) in the complete dataset of 1,955 sequences (light blue, all) grouped according to taxonomy at the phylum (light green) and class level (light yellow); the “other” category (light grey) includes sequences from phyla/classes with <2% of all 1,955 sequences and in which major pilins are not enriched. Grey dots, major pilins ≤165 aa; purple dots, large major pilins (≥166 aa), and blue dots, major pilins of solved T4P of the K02650 group (PilE from *N. meningitidis* (Nm PilE; 5KUA)^22^; PilA from *P. aeruginosa* PAK (Pa PilA; 5VXY)^21^; PilE from *N. gonorrhoeae* (Ng PilE; 5VXX)^21^; PilA4 from *T. thermophilus* (Tt PilA4; 6XXD)^26^; PilA-N from *G. sulfurreducens* (Gs PilA-N; 6VK9)^19^). Error bars, mean length ± standard deviation (STDEV). Black numbers (n) above the x-axis, total number of sequences in a phylum/class. Purple numbers (n), number of large major pilins, and their % of the total number in a phylum/class. Please note that PpdD from enterohemorrhagic *E. coli* (Ec PpdD; 6GV9)^23^, PulG from *K. oxytoca* (Ko PulG; 5WDA)^25^ and PilA5 from *T. thermophilus* (Tt PilA5; 6XXE)^26^ are not depicted. PpdD belongs to K02682 (prepilin peptidase dependent protein D), which is composed of predominantly enterobacterial PpdD homologous sequences with a full length <150aa, PulG belongs to K02456 (GspG, general secretion pathway protein G) and PilA5 is not yet assigned to a KEGG group. **B.** Consensus sequence (con. aa) and consensus secondary structure (con. ss-α) of α1-helix based on 1,955 mature major pilin sequences and major pilins of previously solved T4P structures. The sequences comprising the α1-helix of major pilins of the solved T4P structures as in **A** are included for comparison. For comparison, the sequence comprising the α1-helix of PilA from *M. xanthus* (Mx PilA) is also included^11^. **C.** Schematic of overall architecture of major pilins of solved T4P structures. Red boxes, α1-N and α1-C, with the melted stretch in-between in light red. Numbers, total length of the α1-helix, and the end as well as start of α1-N and α1-C. The sequences used are as in **A**. Note that the major pilin of G. *sulfurreducens* is heterodimeric and composed of PilA-N (61 aa) and PilA-C (104 aa)^19^.

To understand the size variation among the mature major pilins, we performed sequence analyses and secondary structure predictions based on a multiple sequence alignment of the 1,955 major pilins and the major pilins of previously solved T4P structures (Fig. 1B, C). The secondary structure consensus revealed an N-terminal α1-helix with an average length of 51 aa (Fig. 1B), in agreement with a previous study^4^. The amino acid consensus revealed that the N-terminal portion is predominantly hydrophobic and more highly conserved than the C-terminal portion of the α1-helix. The N-terminal portion of α1-helix is inserted in the IM before pilus assembly, which accounts for the hydrophobicity. Comparison of these two regions to the major pilins of solved T4P structures shows that they correspond well to the hydrophobic α1-N and the amphipathic α1-C (Fig. 1B, C). Thus, the size difference among major pilins arises from size differences in the globular domain.

### Cryo-EM structure of the M. xanthus T4P reveals unusual packing

To understand the properties of T4P built from a large major pilin, we focused on T4P^Mx^. The mature PilA (MXAN_5783) has a length of 208 aa with a predicted α1-N highly similar in sequence to the major pilins of the solved T4P structures, but with the predicted amphipathic α1-C containing more basic and less hydrophobic residues than those other major pilins (Fig. 1B). For structure determination of T4P^Mx^, we purified T4P from the hyper-piliated Δ*pilT* strain, in which T4P are extended but not retracted (Fig. 2A, S2A, B). We used cryo-EM to determine the structure of the T4P^Mx^ and obtained the structure at 3.0 Å resolution, the highest resolution so far reported for a T4P structure (Fig. 2B, S2C).

**Figure 2.**
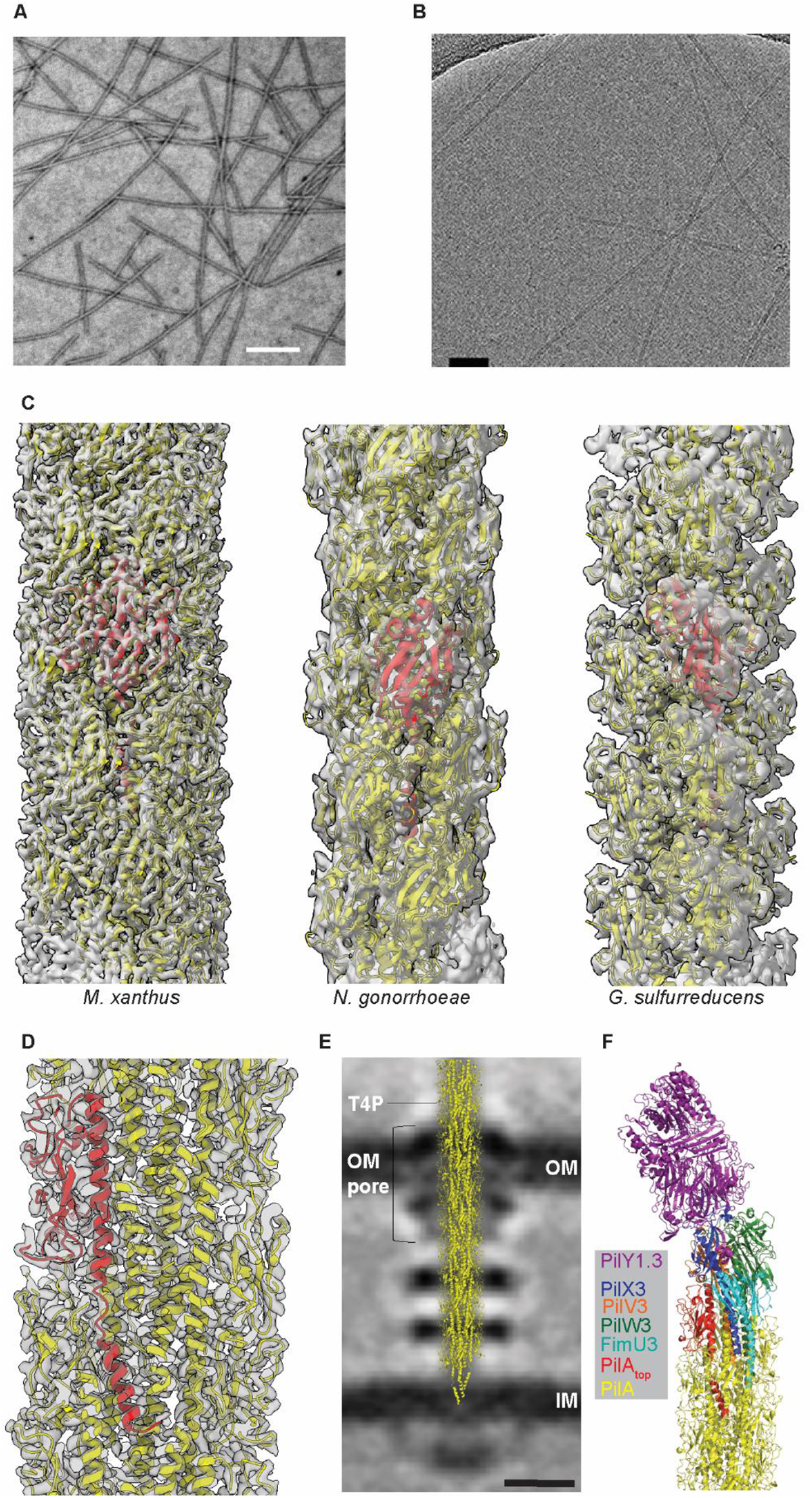
Cryo-EM structure of the T4P^Mx^. **A.** Representative micrograph of negatively stained T4P^Mx^. Scale bar, 200nm. **B.** Representative cryo-EM micrograph of T4P^Mx^. Scale bar, 500nm. **C.** Comparison of cryo-EM reconstructions of T4P and ribbon models of pilins from *M. xanthus* (left), *N. gonorrhoeae* (center), and the 2-chain G. *sulfurreducens* (right), where the transparent surfaces are the cryo-EM density maps, and one pilin subunit is shown in red in each. **D.** Cross section of the cryo-EM reconstruction of the T4P^Mx^ with a single PilA subunit shown in red. **E.** Placement of the T4P^Mx^ structure into the central slice of the subtomogram average of the piliated *M. xanthus* T4PM revealed by cryo-electron tomography^10^. The T4P, the outer membrane pore (OM pore), OM, and IM are indicated. Scale bar, 10nm. **F.** Complete T4P^Mx^ structure composed of T4P^Mx^ and a tip complex. The top PilA of the T4P^Mx^ is shown in red and PilA subunits of T4P^Mx^ below are shown in yellow, the AlphaFold-Multimer model containing the four minor pilins (FimU3, PilV3, PilW3, PilX3) and PilY1.3 is shown in the indicated colors. The complete T4P^Mx^ structure was generated by superposing the top PilA (red) of the T4P^Mx^ with the bottom PilA (wheat) of the AlphaFold-Multimer model (RMSD=0.841, Fig.S2E). The superposed PilA from the AlphaFold-Multimer model is not shown in the complete T4P^Mx^ structure.

The subunits in the filament are related to each other by an azimuthal rotation of 100.7° and an axial rise per subunit of 10.0 Å, generating a right-handed 1-start helix with a pitch of ∼36 Å and 3.6 subunits per turn. The filaments are ∼7 nm in diameter and, in contrast to all previous T4P structures, resemble a rather solid cylinder without the modulation of the surface due to smaller C-terminal domains (Fig. 2C, S3A). The individual PilA subunits within the cryo-EM reconstruction follow the overall blueprint of major pilins in solved T4P structures with the N-terminal α1 generating the core of the pilus and the globular C-terminal domain decorating the surface (Fig. 2D). We also note that the solved structure of the T4P^Mx^ with its diameter of ∼7 nm readily fits into the overall architecture of the *M. xanthus* T4PM, which we previously solved using cryo-electron tomography^10^ (Fig. 2E).

The 3.0 Å-resolution of the T4P^Mx^ structure allowed for building *de novo* an atomic model of individual PilA subunits (Fig. 2D, Fig. 3A). The N-terminal α1-helix of PilA^Mx^ extends from residues 3-54 and contains the hydrophobic α1-N (aa 3-18) and the amphipathic α1-C (aa 24-54) separated by an unfolded stretch of five residues around the conserved P22 (Fig. 1B, 2D, 3A, B, S3B). This is similar to the local melting of this helix seen in major pilins in previous T4P structures^20^ (Fig. 1C, S3B).

**Figure 3.**
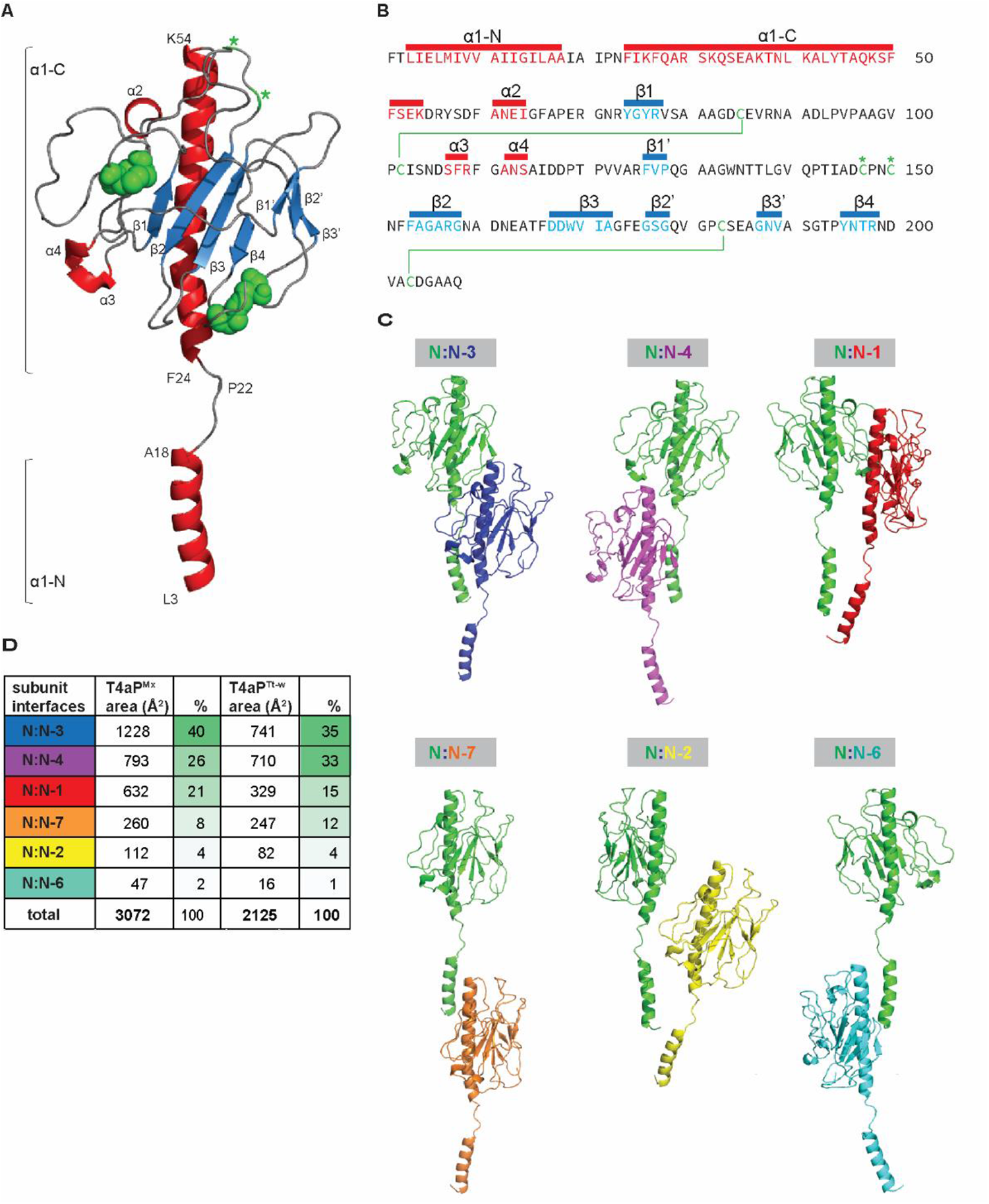
Atomic model of PilA^Mx^ and subunit interface analysis within the T4P^Mx^ filament. **A.** Ribbon representation of PilA^Mx^, with helical elements (α1-4) shown in red, β-stranded elements (β 1-4, β1’-3’) shown in blue, less-structured areas in grey, disulfide bridges shown in green and free cysteines shown with green stars. The first and last residues of the helices of α1-N and α1-C are shown as well as the localization of the conserved Pro 22 within the melted region. **B.** Sequence of PilA depicting structural elements, disulfide bridges, and free cysteines as in **A**. **C.** Ribbon representation of the six different subunit interfaces with the largest three on top and the three smaller ones at the bottom. The color code of the subunits is shown above. **D.** Areas of the six different subunit interfaces of T4P^Mx^ in comparison to the interfaces in the wide T4P of *T. thermophilus* and percentage of those different areas of the total sum shown with a green to white color scale. The color code of the subunit interface corresponds to the subunit interacting with N as in **C**.

The large globular domain (aa 55-208) contains two antiparallel β-sheets, one four-stranded sheet composed of β1-4 and one three-stranded sheet composed of β1’-3’, as well as three α-helices (α2-4) (Fig. 3A, B). These regular structural elements (α-helices, β-strands), which are interrupted by loops, account for ∼26% of the globular domain (Fig. S3B). Compared to the major pilins of the other solved structures, the extra residues in PilA^Mx^ are largely found in a region between β1 and α3 and a region forming β3, β2’ and β3’ (Fig. S3C), and while those previous structures all have a contiguous antiparallel four-stranded β-sheet^19, 21–23, 25–27^, the two antiparallel β-sheets in PilA^Mx^ are non-contiguous (Fig. 3A, B, S3B, C).

Two disulfide bridges are present in the large globular domain. C95/C102 connects and likely stabilizes the region between β1 and α3. C183/C203 connects the β’-sheet to the C-terminal part of the globular domain (Fig. 3A, B), somewhat similar to the C-terminal D-region known from other pilins that attaches the β-sheet to the C-terminal portion of the globular domain^4^.

### The large globular domain is involved in extensive intersubunit interactions

Within the T4P^Mx^ the globular domain of an individual PilA monomer extensively interacts with neighboring subunits (Fig. 3C), including six different subunit-subunit interfaces, three large (N:N-3, N:N-4, N:N-1) and three small (N:N-7, N:N-2, N:N-6) (Fig. 3C, D). In total, these interfaces add up to ∼ 3000 Å^2^ of buried surface area (Fig. 3D). Because a pilin subunit interacts with pilins above and below, every individual pilin subunit has a total of 12 interaction partners, adding up to a total of ∼6000 Å^2^ of buried surface area per pilin.

The structure of the wide T4P of *T. thermophilus (*T4P^Tt-w^) composed of the 125 aa PilA4 pilin was also solved at a high resolution (Fig. S3A)^26^, allowing a direct comparison of subunit interface areas between the T4P^Mx^ and T4P^Tt-w^. T4P^Tt-w^ also has six subunit interfaces and similarly to the T4P^Mx^, the largest interfaces occur between N:N-3, N:N-4 and N:N-1 (Fig. 3D). In comparison to the T4P^Tt-w^, there is a ∼50% increase in the buried interfacial area per subunit in the T4P^Mx^ (Fig. 3D), deriving largely from more extensive interactions in the N:N-3 and N:N-1 interfaces. Similarly, a comparison of the N:N-3, N:N-4, and N:N-1 interfaces of T4P^Mx^ with those of the lower resolution T4P structures of the *E. coli* EHEC (major pilin, 140aa), *N. gonorrhoeae* (major pilin, 158aa), *N. meningitidis* (major pilin, 161aa), and *P. aeruginosa* PAK (major pilin, 143aa) shows that these three interfaces in these four structures vary from ∼1500-2000 Å^2 23^, and are thus also significantly smaller than in the T4P^Mx^.

### A structural model of the complete T4P^Mx^ including the tip complex

*M. xanthus* encodes three sets of each four minor pilins and one PilY1 adhesin^11, 14, 15^. At least two of these three sets, i.e. those encoded by gene cluster_1 and gene cluster_3, not only form a priming complex for pilus assembly but also a tip complex involved in adhesion^11, 14, 15^. Similar to major pilins, minor pilins are composed of an N-terminal α1-helix and a globular C-terminal domain^4^; PilY1 proteins share a conserved C-terminal domain while the N-terminal domain is more variable^43^.

For the complex formed by the cluster_3 proteins, it was proposed that the less conserved N-terminal domain of PilY1.3 sits at the top while the conserved C-terminal domain interacts with a complex composed of four minor pilins below, which, in turn, interact with PilA below^11^. Specifically, based on pull-down experiments and direct interaction analyses, the minor pilin PilX3 was placed directly below PilY1.3, followed by PilW3, FimU3, PilV3 and PilA ^11^. To generate the first complete structural model of a T4P, we first generated a structural model using AlphaFold-Multimer of the tip complex composed of one copy each of the major pilin, the four minor pilins and PilY1 using the proteins of cluster_3^11^.

A high confidence AlphaFold-Multimer model (Fig. S2D), largely confirmed the suggested organization of this complex with PilY1.3 at the top followed by PilX3, PilV3, PilW3, FimU3 and PilA at the base, i.e. the only difference is the placement of PilV3 between PilW3 and PilX3 (Fig. S2E). In the AlphaFold-Multimer model only α1 of PilA is slightly kinked (around the conserved P22) (Fig. S2E), while the four minor pilins lack that residue^11^. Interestingly, a stretch of eight amino acids residues of the C-terminal end of PilY1.3 is modeled to form a β-strand, which, together with two β-strands of PilX3, forms an three-stranded antiparallel β-sheet (Fig. S2F), suggesting that PilX3 and PilY1.3 interact by β-strand addition, more precisely by β-sheet augmentation^44^. Interestingly, despite substantial sequence diversity between cluster_3 and cluster_1 components^11, 14^, the same order of components as well as the proposed β-sheet augmentation between PilX1 and PilY1.1 is predicted for the high confidence AlphaFold-Multimer model of the cluster_1 proteins (Fig. S2G, H, I). Protein-protein interactions by β-strand addition are also involved in the assembly and stabilization of the Type 1 pilus^45^, and are reported to be extraordinarily stable against dissociation and unfolding^46^. To generate the complete model of the T4P^Mx^ including the tip complex, we fitted the model of the tip complex of the cluster_3 proteins into the T4P^Mx^ structure by superposing the top PilA of the T4P^Mx^ with the PilA of the AlphaFold-Multimer model (Fig. 2F). Importantly, these two PilA molecules could readily be superposed giving rise to a structure in which the four minor pilins tops the T4P^Mx^ followed by the PilY1.3 adhesin firmly attached through its C-terminal domain to PilX3 via β-sheet augmentation.

### T4P^Mx^ has increased bending and axial stiffness compared to less compact T4P

Since the resistance to bending will scale as the fourth power of the radial mass distribution, we expected that the increased contacts between the outer domains near the outside of the pilus would make the T4P^Mx^ filament more rigid than previously studied ones. We quantified its bending stiffness with the persistence length (PL). Because the persistence length is derived from an analysis of fluctuations in curvature from filaments at thermodynamic equilibrium, cryo-EM is ill-suited for making such measurements, due to the large forces present from both fluid flow during blotting and the compression of long filaments into a thin film^47, 48^. Consequently, we used purified T4P^Mx^ visualized by negative stain transmission electron microscopy (TEM) and determined a PL of 21µm for T4P^Mx^ (Fig. S3A, S4A). In parallel experiments, we determined the PL of the less compact T4P of *N. gonorrhoeae* (major pilin, 158aa; Fig. S3A) and *P. aeruginosa* PAK (major pilin, 144aa; Fig. S3A) as 11µm and 13µm, respectively (Fig. S3A, S4A). We conclude that the more extensive C-terminal domain contacts in the T4P^Mx^ do indeed result in increased bending stiffness.

We also expected that these increased contacts would reduce the axial compliance, resulting in a greater force needed to extend these filaments. We therefore analyzed the force-extension behaviour and adhesive properties of T4P^Mx^ in live cells using atomic force microscopy (AFM) force spectroscopy (FS) as described for T4P in *P. aeruginosa* PAO1 and PA14 (major pilin length:143 & 173 aa, respectively)^28, 30, 49^. Those studies reported two distinct force-extension profiles when single T4P on live cells were pulled: (i) a tensile force that initially increased in an approximately linear fashion with pilus stretching before rupturing of the contact between the pilus and the AFM tip; and (ii) an initial increase in force followed by a constant force plateau before rupture occurred. Because of the approximate linearity of the first type, these profiles were called linear nanosprings, and their spring constant *k_pilus_*, which is a measure of pilus axial stiffness, was quantified as ∼2 pN/nm^28, 30^. We applied the AFM-FS methodology used in^30^ to characterize T4P^Mx^ on live cells (Methods). Briefly, we covalently modified a gold AFM tip to make it hydrophobic. Subsequently, single *M. xanthus* cells adhering to a polystyrene surface were visualized with an inverted microscope and force probed in buffer with the hydrophobic AFM tip. Specifically, because T4P^Mx^ are localized to one of the cell poles^50^, the AFM probe, initially at a specified height above the sample, is displaced downwards close to a piliated cell pole until a T4P, which is freely moving in the buffer, by chance adheres via hydrophobic interactions to the AFM tip (Fig. 4A, i). Then the probe is moved upwards at a constant velocity, thereby also lifting the relaxed pilus (Fig. 4A, ii) until it reaches its initial height (Fig. 4A, iii). During this movement, a bound T4P is loaded with tension causing its extension and resulting in the downward bending of the cantilever, thereby allowing the quantification of the tensile force (Fig. 4A, iii). Once the tensile force exceeds the strength of the interactions between the pilus and the AFM tip, the contact between the pilus and the AFM tip ruptures and the cantilever relaxes (Fig. 4A, iv). The T4P extension until rupture of the pilus-AFM tip contact is recorded as force-distance (*F*-*d*) curves (Fig. 4A). From such *F*-*d* curves over a raster grid in force volume mode (Methods), we constructed correlated topographic and adhesion maps, thereby pinpointing the exact location of pilus signatures (Fig. S4B). As previously observed for *P. aeruginosa* T4P, we observed both nanospring (Fig. 4B) and force plateau signatures (Fig. S4C) almost exclusively close to one of the cell poles for wild-type (WT) *M. xanthus* cells. By contrast, such signatures were nearly absent in *M. xanthus* Δ*pilA* cells (Fig. S4D). The measured rupture forces in the pilus nanosprings (∼120pN) (Fig. 4C) and force plateaus (∼220pN) (Fig. S4C) were similar to those reported for *P. aeruginosa* T4P^28, 30, 49^, indicating no differences in the adhesive properties between *M. xanthus* and *P. aeruginosa* pili. The spring constant *k_pilus_* of T4P^Mx^ at low (1µm/s as in^30^) and at moderate (5µm/s as in^28, 49^) pulling speeds was ∼4.0pN/nm and ∼5.5pN/nm, respectively (Fig. 4C). Importantly, these values are at least two-fold higher than those reported for T4P in *P. aeruginosa* PAO1 and PA14^28, 30^, indicating a greater average axial stiffness of the T4P^Mx^ and that T4P^Mx^ are more resistant to stretching than T4P of *P. aeruginosa*.

**Figure 4.**
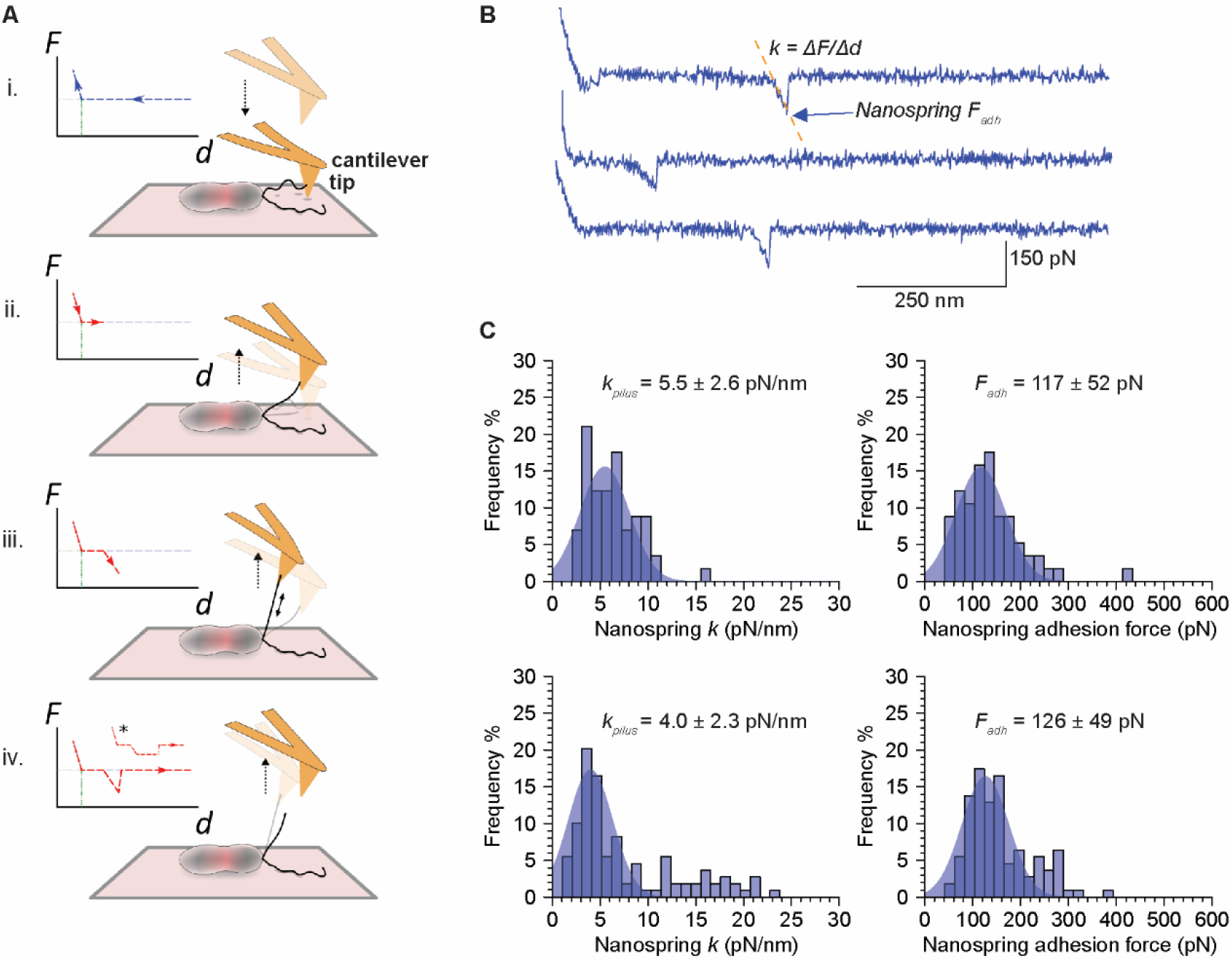
T4P^Mx^ have increased axial stiffness and elastic properties and undergo force-induced elongation in response to pulling forces by AFM-FS. **A.** Schematic of AFM-FS approach to collect force-extension profiles of *M. xanthus* T4P. (i) Cells in buffer were adhered to a polystyrene surface. The hydrophobic AFM probe on the cantilever approached towards the surface near one of the cell poles until it touches and pushes on the surface of the dish or cell, causing the upwards bending of the cantilever that is recorded as a positive force in the approach Force-distance (*F*-*d*) curve (blue). By chance, a T4P freely floating in the buffer adheres to the hydrophobic tip (i.e. hydrophobic contacts are established between the solvent accessible surface of the pilus fiber and the CH_3_ groups exposed on the AFM tip). (ii) The AFM tip is retracted away from the surface lifting the relaxed T4P until (iii) the T4P becomes loaded with tension resulting in downwards bending of the cantilever, thereby allowing quantification of the tensile force (red *F*-*d* curve). (iv) When the tensile force within the T4P exceeds that of the bonds between the pilus and the hydrophobic AFM tip, the contact between the two is ruptured and the cantilever relaxes resulting in a zero-force measurement (baseline) in the *F*-*d* curve. Two types of signatures are expected for T4P, linear (Hookean) nanosprings and constant force plateaus (indicated with an asterisk in the *F*-*d* curve). **B.** Three representative *F*-*d* curves obtained for WT *M. xanthus* cell showing nanospring signatures. The rupture/adhesion force (*F_adh_*) is indicated by a blue arrow, the dotted line indicates the slope of the nanospring profile used to determine its spring constant (*k_pilus_*). **C.** Histograms showing the distribution of nanosprings *k_pilus_* (left) and *F*_adh_ (right) as determined from *F*-*d* curves generated at a probe retraction velocity of 1µm/sec (top, n= 57 nanosprings in 82/1312 curves from 5 tip-cell combinations) or 5µm/sec (bottom, n= 109 nanosprings in 44/5400 curves from 5 tip-cell combinations). Numbers indicate mean ± STDEV.

We also note that even though the mean rupture forces in the pilus nanosprings (Fig. 4C) and force plateaus (Fig. S4C) were in close agreement with the reported values for *P. aeruginosa* T4P, T4P^Mx^ can resist pulling forces up to 400-500 pN before the contact between the pilus and the AFM tip ruptures (Fig. S4C) while T4P of *P. aeruginosa* resisted forces only up to 250 pN^30^. Nevertheless, T4P^Mx^ have elastic properties and undergo a force-induced elongation in response to pulling forces as previously described for T4P of *N. gonorrhoeae, N. meningitidis* and *P. aeruginosa*^28–31^ consistent with the previous hypothesis that the extensibility arises from further melting of the N-terminal α-helix^21^.

### Disruption of PilA subunit-subunit interfaces reduce persistence length

Among the six interfaces between PilA subunits in the T4P^Mx^, N:N-3, N:N-4 and N:N-1 are not only the largest contributors to the subunit interface but also significantly more extensive than those in the T4P^Tt-w^ (Fig. 3C-D), and the T4P of *E. coli* EHEC, *N. gonorrhoeae*, *N. meningitidis,* and *P. aeruginosa* PAK. To assess how these three interfaces contribute to pilus bending stiffness and to T4P function *in vivo*, we mutagenized charged residues engaged in salt bridge formation in these three interfaces (Fig. 5A-C). Specifically, we targeted the residues R30, K37, E53 at the N:N-3 interface, D55, R73, R109 at the N:N-4 interface, and K48, E69, R70 at the N:N-1 interface and substituted these residues separately with residues with either a polar side chain (Asn or Gln) or Ala. These nine residues are either localized in α1-C (R30, K37, K48, E53) or in the globular domain (D55, E69, R70, R73, R109) (Fig. 3B), and are forming salt bridges connecting either α1-helices (R30, K37, E53 at the N:N-3 interface), globular domains (D55, R73, R109 at the N:N-4 interface, R70, E175 at the N:N-1 interface), or α1-helices and globular domains (K48, E69 at the N:N-1) of the corresponding subunits (Fig. 5A-C). The corresponding 18 mutations were introduced into the *pilA* gene at the native locus of the WT and in the retraction-deficient Δ*pilT* mutant to distinguish between T4P extension and hyper-retraction defects caused by these substitutions.

**Figure 5.**
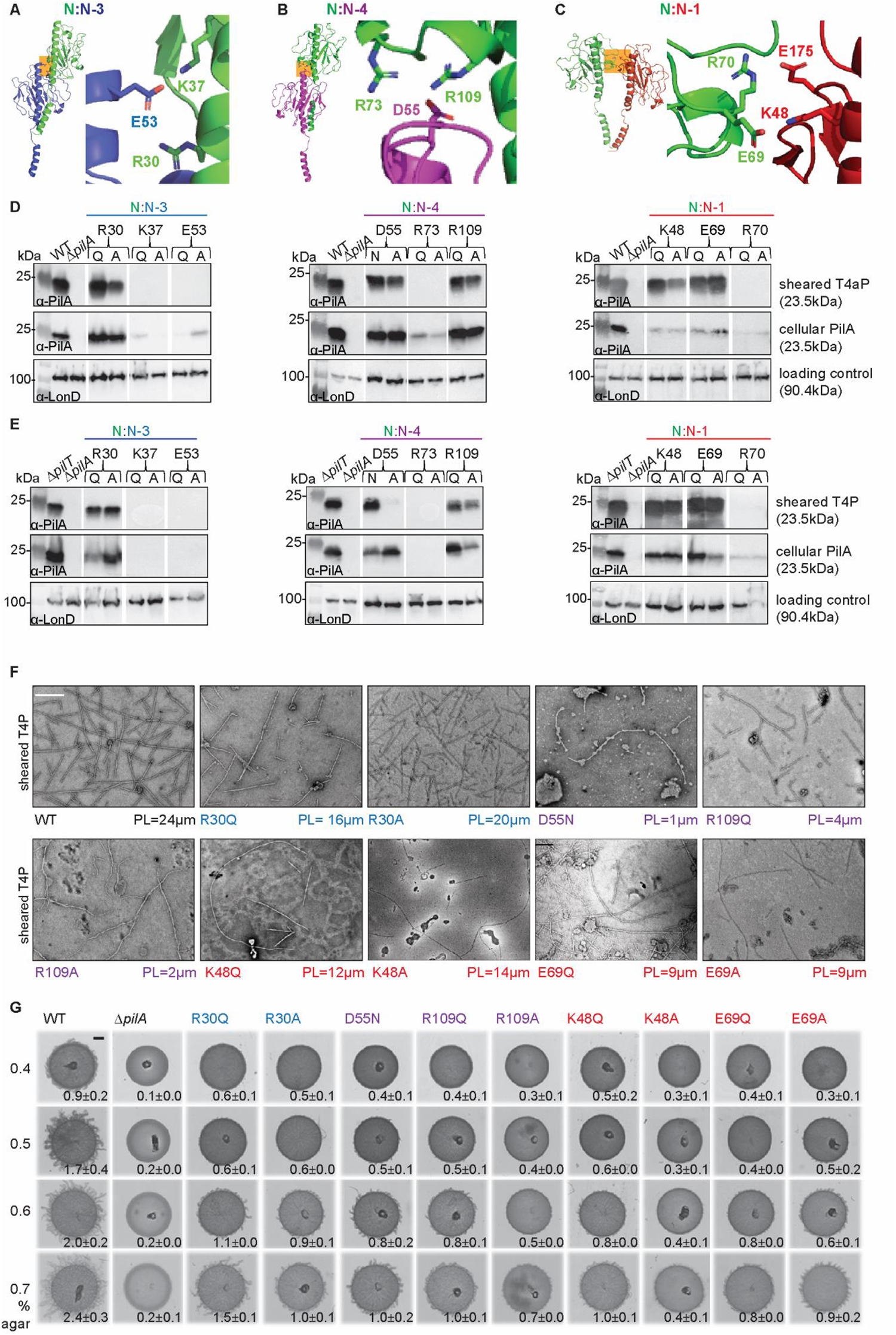
Mutagenesis of selected charged residues in PilA subunit interfaces. **A-C.** Ribbon representation of interacting subunits N:N-3 (A), N:N-4 (B) and N:N-1 (C) as overviews (left parts) with orange boxes indicating the position of the magnified views (right parts) depicting the charged residues in these areas. The depicted residues, except for E175, were all targeted for mutagenesis. The subunit color code is as in Fig. 3C. **D-E.** Effect of amino acid substitutions on PilA* accumulation and T4P formation. T4P sheared off from the surface of cells (top rows), and total cell extracts (middle rows) were separated by SDS-PAGE and probed with α-PilA antibodies. The membrane used for cell extracts was stripped, and probed with α-LonD antibodies as a loading control (bottom rows). Material from the same amount of cells was loaded per lane. The PilA variants were analyzed in WT (**D**) and the Δ*pilT* mutant (**E**). Proteins with their calculated molecular masses and positions of molecular markers are indicated. Gaps between lanes indicate lanes that were deleted for presentation purposes. **F.** Negative staining micrographs of sheared T4P. PL values in µm are shown below the images. Scale bar, 250 nm. Note that these pili, in contrast to those in Fig. 2A, were imaged directly after the precipitation and not further purified by sucrose gradient centrifugation. **G.** Substitutions at intersubunit interfaces perturb T4P-dependent motility. Cells were incubated 24 hrs before imaging. The Δ*pilA* mutant served as a negative control. Scale bar, 1 mm. Numbers indicate mean±STDEV of increase in colony diameter in 24hrs from two independent experiments.

We first examined the accumulation of the PilA variants in total cell extracts and their ability to support T4P formation. Substitutions of four of the nine residues (K37, E53 of α1-C and R70, R73 of the globular domain) (Fig. 5A-C) caused strongly reduced or abolished PilA* accumulation in total cell extracts of both strain backgrounds (Fig. 5D, E). Consistently, in a T4P shear-off assay in which pili are sheared off the surface of cells, these PilA variants did not support T4P formation in either strain background (Fig. 5D, E). Thus, these residues are important for PilA stability and lack of T4P formation is neither due to an extension defect nor a hyper-retraction defect.

Mutagenesis of the remaining five residues allowed PilA* accumulation in total cell extracts in both strain backgrounds at the same or slightly lower level than PilA^WT^ (Fig. 5D, E). Except for the D55A variant, they supported T4P formation in both strain backgrounds at essentially the same level as PilA^WT^ (Fig. 5D, E). Paradoxically, the D55A variant, while accumulating in both strain backgrounds, only supported T4P formation in the WT but not in the Δ*pilT* mutant. From here on, we focused on the nine variants that accumulated and supported T4P formation in both strain backgrounds.

To assess the mechanical properties of the nine variant T4P, we purified them from the Δ*pilT* background and determined their PL (Fig. 5F). All nine variants had a moderately to strongly reduced PL (Fig. 5F, S5). In particular, substitutions in the N:N-4 interface caused dramatic reductions in PL (Fig. 5F), while substitutions in the N:N-3 and N:N-1 interfaces generally only caused a ∼50% reduction in PL (Fig. 5F). We conclude that the substitutions do not interfere with the extension of T4P; however, the T4P assembled by the PilA variants have decreased bending stiffness.

### Disruption of PilA subunit-subunit interfaces reduces T4P-dependent motility

To analyze whether PilA subunit interface disruption affects T4P-dependent motility, we analyzed the *M. xanthus pilT*^+^ strains synthesising these nine variants. *M. xanthus* moves by T4P-dependent motility, which is favoured on soft, moist surfaces, and by gliding motility, which is favoured on hard agar^36^. Surface stiffness was reported to stimulate T4P-dependent motility in *P. aeruginosa*^51^. Therefore, we tested WT as well as the strains expressing the nine PilA variants on soft agar of different stiffness by using a range of agar concentrations (0.4-0.7%) and the increase in colony diameter at 24 hrs as a readout for T4P-dependent motility. The Δ*pilA* strain, which only moves by gliding motility, served as a negative control for T4P-dependent motility and to verify that gliding motility did not significantly contribute to the increase in colony diameter under these conditions.

The WT displayed T4P-dependent motility on all four agar surfaces generating the characteristic flares at the colony edge, and the colony diameter increased ∼2.5-fold with the agar concentration, while the Δ*pilA* mutant, as expected, generated smooth-edged colonies and only displayed a minor increase in colony diameter (Fig. 5G). These findings are consistent with the observation that surface stiffness stimulates T4P-dependent motility in *P. aeruginosa*. All strains expressing a PilA variant had strongly reduced T4P-dependent motility at all agar concentrations (Fig. 5G). Like the WT, they generally showed improved T4P-dependent motility with increasing agar concentrations; however, none reached the WT level even at 0.7% agar (Fig. 5G). We conclude that WT T4P^Mx^ supports T4P-dependent motility on surfaces of different stiffnesses and more efficiently at higher agar concentrations. In contrast, the T4P of the variants are less efficient at supporting motility under all the tested conditions and are only slightly stimulated on stiffer surfaces.

## Discussion

Here, we elucidate the structure of the T4P^Mx^ using cryo-EM at a resolution of 3.0 Å and demonstrate that, in contrast with all previous T4P structures, the T4P^Mx^ structure is highly compact. The PL of T4P^Mx^ is ∼2-fold higher than those of *P. aeruginosa* and *N. gonorrhoeae*, consistent with the greatly increased contacts at higher radius in the T4P^Mx^. Similarly, the spring constants of T4P^Mx^ at low and moderate pulling speeds are at least 2-fold higher than those reported for *P. aeruginosa* T4P, indicating a greater axial stiffness of T4P^Mx^. Also, T4P^Mx^ can resist pulling forces up to 400-500 pN, in agreement with the observation that T4P^Mx^ can resist forces up to 150pN generated during retractions^7^. These data make the T4P^Mx^ the strongest and most rigid T4P yet described. Nevertheless, T4P^Mx^ have elastic properties and undergo a force-induced elongation in response to pulling forces as previously described for T4P of *N. gonorrhoeae, N. meningitidis,* and *P. aeruginosa*^28–31^.

The T4P^Mx^ is more compact, more rigid, and stronger than other T4P due to the larger globular domains, which are involved in more extensive intermolecular interactions than seen in other T4P structures. The larger C-terminal globular domains provide surfaces for extensive interactions, causing a measurable increase in total interface area, and allow every individual pilin to interact with six pilins above and six below. Among these six interfaces, the three largest are the N:N-3, N:N-4, and N:N-1, and all three interfaces contain residues that engage in intersubunit salt bridges, i.e. R30_N_ and/or K37_N_ and E53_N-3_ in N:N-3, R73_N_ and/or R109_N_ and D55_N-4_ in N:N-4, and E69_N_ and K48_N-1_, R70_N_ and E175_N-1_in N:N-1 (Fig. 5A-C). Disruption of these three subunit interfaces reduced the bending stiffness of the corresponding T4P^Mx^ variants but the disruption of the salt bridge connecting two globular domains in the N:N-4 interface had the strongest effect, supporting the hypothesis that the large globular domains contribute significantly to the increased bending stiffness.

We also observed that the PilA variants K37Q/A, E53Q/A, R70Q/A, R73Q/A had reduced stability (Fig. 5D, E), suggesting that these residues are important for intramolecular interactions before the incorporation of PilA into the T4P and that mutagenesis of these residues causes misfolding and degradation of PilA*. This is similar to earlier findings, showing that mutagenesis of A18, I19 and A20 of PilA^Mx^ (I19 and A20 are part of the melted region, Fig. 3A, B) can strongly affect PilA accumulation^52, 53^. Interestingly, mutagenesis of other major pilins also support the notion that intra-and intersubunit salt bridges contribute to the stability of the pilin, assembly of the T4P, and T4P function^21, 23, 25, 54–56^.

T4P-dependent motility was stimulated by increased substrate stiffness. A similar observation was made in *P. aeruginosa,* and it was suggested that this stimulation involves an increased probability of T4P retraction on the stiffer agar surface^51^. Interestingly, the PilA variants in which subunit interfaces were disrupted supported T4P extension as efficiently as native PilA. However, the T4P made from these PilA variants were less efficient at supporting T4P-dependent motility than PilA^WT^ at all substrate stiffnesses. The T4P made from the PilA variants had a reduced PL, indicating decreased bending stiffness or flexural rigidity. However, the PL did not correlate with the ability to support motility suggesting that it is not the decreased bending stiffness *per se* that results in the motility defect. Also, even variant T4P with PLs similar to those of T4P of *P. aeruginosa* and *N. gonorrhoeae* did not support motility. During the extension/adhesion/retraction cycles, only retractions generate a force sufficient to pull a cell forward^5, 6^, suggesting that the variant T4P likely have retraction defects. The hexameric PilB and PilT ATPases that power extension and retraction, respectively, bind at the base of the T4PM in a mutually exclusive manner^10^. The swap from PilB to PilT, and thus initiation of retraction, was suggested to be a stochastic event^57^, or, alternatively, it was suggested that it is induced by adhesion of the pilus tip to the substratum in a process in which tip adhesion causes conformational changes in the pilus that are communicated to the base of the T4PM^58^. *N. gonorrhoeae* and *P. aeruginosa* T4P, as also reported here for T4P^Mx^, undergo force-induced conformational changes to elongate^28–30^. Therefore, we speculate that the motility defect of the T4P variants could be caused by (1) less efficient transmission of conformational changes from the tip to the base of the T4PM to stimulate the swap from PilB to PilT, (2) reduced ability to undergo force-induced conformational changes during retraction, or (3) even breakage of the T4P when it is pulled taut during retraction.

Our results advance our understanding of how sequence divergence of major pilins shapes the functional properties of T4P. Moreover, the information gained from the first complete T4P model, composed of major pilins, four minor pilins, and a large PilY1 adhesin, provides insights into the interactions between major and minor pilins, as well as between minor pilins and PilY1.In future studies of T4P, it will not only be interesting to obtain the structure of T4P formed by large major pilins from different bacteria to further reveal their ecological relevance but also to obtain detailed insights into the interactions between the tip complex and the remaining T4P.

## Methods

### Bioinformatics

Sequences of the K02650 (type IV pilus assembly protein PilA) were extracted from the KEGG SSDB database^59^. To filter out highly homologous sequences we used the cdhit program with a threshold of 90% sequence identity^60^. The 2308 obtained sequences were analyzed for presence of T3SP and subsequently processed into the mature pilin form using SignalP (6.0)^61^. The taxonomic classification of the remaining 2071 pilin sequences was collected from KEGG SSDB database and sequences without a bacterial classification as well as sequences from bacterial phyla, only represented by one genome, were excluded from the analysis. The 1955 remaining pilin sequences were analyzed with the PROMALS3D multiple sequence and structure alignment server^62^ to obtain aa and secondary structure consensus sequences on the base of the previously solved T4P structures.

Alignments were generated using T-Coffee^63^ and the ClustalW output format^64^. They were shaded using the BoxShade Server or the BioEdit sequence alignment editor (7.2.5)^65^. For the phylogenetic tree, the ANCESCON tool^66^ of the MPI bioinformatics Toolkit^67^ was used. The phylogenetic tree was annotated using iTol (v6)^68^.

### Bacterial strains, plasmids and growth media

Strains and plasmids are listed in Supplementary Tables 3 and 4, respectively. All *M. xanthus* strains are derivatives of the DK1622 WT strain^50^. In-frame deletion mutants were generated as described^69^. All plasmids were verified by sequencing. All strains were confirmed by PCR and sequencing. Oligonucleotides are listed in Supplementary Table 5. *M. xanthus* strains were grown at 32°C in 1% casitone broth (CTT) (1% casitone, 10mM Tris-HCl pH 8.0, 1mM KPO_4_ pH 7.6, 8mM MgSO_4_) or on 1% CTT, 1.5% agar plates supplemented with kanamycin (40µg ml^-1^) when required^70^. Growth was followed by measuring optical density at 550nm (OD_550_). *E. coli* strains were grown in lysogeny broth (LB)^71^. Plasmids were propagated in *E. coli* Mach1.

### T4P purification and T4P shearing assays

T4P were sheared off from the hyper-piliated Δ*pilT* strain using a protocol based on the procedure of^72^. Briefly, cells grown on 1% CTT, 1.5% agar plates for 2-3 days were gently scraped off the agar and resuspended in 4 ml/plate pili resuspension buffer (100mM Tris-HCl pH 7.6, 150mM NaCl). The pooled suspension was centrifuged for 20 min at 13,000 *g* at 4°C to remove cell debris. The supernatant was centrifuged twice for 10 min at 13,000 *g* at 4°C. T4P in the cell-free supernatant were precipitated by adding 10× pili precipitation buffer (final concentrations: 100mM MgCl_2_, 500mM NaCl, 2% PEG 6000) for at least 2 hrs at 4°C. The solution was centrifuged for 30 min at 13,000 *g* at 4°C, and the pellet resuspended 1ml pili resuspension buffer. The pili solution was loaded on top of a centrifuge tube containing a 10-70% sucrose gradient (29 ml) of pili resuspension buffer. After 15 hrs centrifugation at 115,000xg in a swing bucket rotor (SW72Ti) at 4°C, the tube was punched at the bottom, and 1.5 ml fractions harvested and analyzed by SDS-PAGE using SDS-lysis buffer (10% (v/v) glycerol, 50mM Tris-HCl pH 6.8, 2mM EDTA, 2% (w/v) SDS, 100mM DTT, 0.01% bromphenol blue). To remove the sucrose, PilA-containing fractions were diluted 13.5 fold in pili resuspension buffer, and the solutions precipitated again with pili precipitation buffer (s.a.). The pili were resuspended in pili resuspension buffer.

For T4P shearing assays, 60 mg cells grown on 1% CTT, 1.5% agar plates for 2-3 days were gently scraped off the agar and resuspended in pili resuspension buffer. Cell suspensions were vortexed for 10 min at the highest speed. Cells from a 100 µl aliquot were harvested, the pellet solved in 100 µl SDS lysis buffer, and immediately denatured at 95°C for 5 min. This represents the cellular fraction. The remaining suspension was centrifuged for 20 min at 13,000 *g* at 4°C. The supernatant was removed and centrifuged twice for 10 min at 13,000 *g* at 4°C to remove cell debris. T4P in the cell-free supernatant was precipitated by adding 10× pili precipitation buffer for at least 2 hrs at 4°C. The solution was centrifuged for 30 min at 13,000 *g* at 4°C, and the pellet was resuspended in SDS lysis buffer (1µl per mg vortexed cells). T4P sheared and purified from the same amount of cells were loaded and separated by SDS-PAGE.

### Transmission electron microscopy

Resuspended solutions of sheared T4P were applied on 300 mesh Formvar/carbon copper-grids. After 10 min, grids were washed twice with water and negative-staining was done with a solution based on an Organotungsten compound (Nano-W, Nanoprobes). Grids were inspected with a JEM-1400 electron microscope (JEOL) at 100 kV.

### Cryo-EM sample preparation and data collection

2 μl of a T4P^Mx^ sample was applied to plasma-cleaned lacey carbon grids, followed by plunge-freezing in liquid ethane using a Leica EM GP. Data collection was carried out at liquid nitrogen temperature on a Titan Krios microscope (Thermo Fisher Scientific) operated at an accelerating voltage of 300 kV. Using a K3 camera (Gatan), 40 movie frames were collected with a total dose of ∼55 electron/Å^2^ and sampling of 1.08Å/pixel. The movies were collected with defocus values ranging between -1 to -2.5 μm.

### Cryo-EM image processing and reconstruction

All the subsequent data processing was performed in CryoSPARC. Raw movie frames were used in the motion correction, followed by CTF estimation. Images with poor CTF estimation were eliminated. Filament tracer was used for filament picking and a total of 6,155,222 256 px-long segments were extracted from 13,006 CTF-corrected images. After final round of 2D classification, 1,312,119 segment remained and was subjected to homogenous refinement, which yield a map of recognizable secondary structure features when imposing the 1-start helical symmetry. The helical parameters converged to a twist of 100.7° and a rise of 10Å per subunit. The resolution of the final reconstruction was determined by two-independent half maps showing a resolution of 3.0Å at FSC=0.143.

### Model building and refinement

An initial homologous model generated via SWISS-MODEL was docked into the cryo-EM map by rigid body fitting in Chimera^73^ and manually edited the model in Coot^74^. The modified monomeric model was then real-space refined using Phenix^75^ to improve the stereochemistry as well as the model-map correlation coefficient. The refined monomeric model was re-built by RosettaCM with helical symmetry, followed by another three rounds of real-space refinement to reduce subunit clashes. The refined symmetrical model was validated with MolProbity^76^, and the coordinates were deposited to the Protein Data Bank with the accession code 8TJ2. The corresponding cryo-EM map was deposited in the EMDB with accession code EMD-41298.

### AlphaFold-Multimer model building

Structures were predicted using AlphaFold-Multimer modeling via ColabFold (Version 1.5.0)^77–79^. The predicted Local Distance Difference Test (pLDDT) and predicted Alignment Error (pAE) graphs of the five models generated were made using a custom Matlab R2020a (The MathWorks) script^80^. Per residue model accuracy was estimated based on pLDDT values (>90, high accuracy; 70-90, generally good accuracy; 50-70, low accuracy; <50, should not be interpreted)^78^. Relative domain positions were validated by pAE. The pAE graphs indicate the expected position error at residue X if the predicted and true structures were aligned on residue Y; the lower the pAE value, the higher the accuracy of the relative position of residue pairs and, consequently, the relative position of domains/subunits/proteins^78^. PyMOL version 2.4.1 (Schrödinger LLC) was used to analyze and visualize the models. Structure superposition of the top PilA of the T4P^Mx^ with the bottom PilA of the AlphaFold-Multimer model was done using the align function in PyMol (root mean square deviation=0.841). The protein sequences of the mature minor pilins and PilY1 proteins without their signal peptides as reported earlier^11^, were used for generating the models.

### Determination of subunit interface areas

The interfacial areas were determined by PDBePISA (https://www.ebi.ac.uk/pdbe/pisa/^81^.

### Persistence length determination

To determine the flexibility of different T4P, persistence length measurements were performed using micrographs of negatively stained T4P from indicated strains. For each strain 30-50 filaments were traced using the ImageJ analysis tool^82^. Persistence length (L) is determined by the statistical relationship of cos (θ) and contour length (λ), according to exp(-λ/L)=<cos(θ)>.

### AFM tip functionalization and FS

Gold AFM probes (PNP-TR, NanoWorld) were incubated overnight in a 1mM ethanolic 1-dodecanethiol solution to modify them with methyl groups and render them hydrophobic, then rinsed with ethanol and sterile water and kept in milliQ water at 4°C until use (no longer than 48 hrs). Early exponential-phase *M. xanthus* cultures (OD_550_ ∼ 0.5) grown in CTT at 32°C in the dark, were diluted in MC7 buffer (10 mM MOPS pH 7.6, 1 mM CaCl_2_) and passaged gently through a 26 gauge needle to dissolve cell aggregates before seeding cells in a 35mm untreated polystyrene Petri dish (Corning). After 30 min incubation to allow cells to adhere, they were adequately adhering for high-quality AFM-FS, gently rinsed with MC7 buffer and immediately used for AFM. Prior to any AFM measurements, the spring constant of the probe’s cantilever was determined as reported previously^83^ allowing for the accurate correlation between measured cantilever deflection with tensile force in stretched T4P. AFM recordings were done at room temperature using a NanoWizard® 4 NanoScience AFM (JPK Instruments) in force mapping (volume) mode, which allows the recording of *F*-*d* curves in a pixel-by-pixel fashion over a defined surface area in a raster array. Sample height could be determined from the approach section of each *F*-*d* curve, while the retract portion provided pilus forced extension and adhesive information and hence correlated images of sample topography and pilus mechanical properties allowing mapping of pilus signatures with respect to the cell body. For each tip-cell pair, a large (10×10 µm, 32 × 32 pixels) force map was first recorded to generate a topographical image of a whole cell and to find pilus signatures. Subsequently, a smaller (3×3 µm, 32×32 or 16×32 pixels) map was recorded over a piliated pole. Cells were visualized using an inverted microscope prior to commencement of AFM force mapping. *F*-*d* curve analysis was performed using the JPK data processing software. The nanospring constant (given as *k_pilus_* below) was calculated using the serial spring equation: 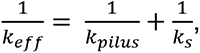 with *k_eff_* (effective spring constant) equal to the slope of the linear region of a nanospring extension profile in an *F*-*d* curve and *k_s_* (spring constant of the cantilever sensor). Graphs pertaining to AFM data was generated using R Studio.

### T4P-dependent motility assays

Cells from exponentially growing *M. xanthus* cultures were harvested and resuspended in 1% CTT to a calculated density of 7×10^9^ cells ml^-1^. 5µl of cell suspension were spotted on soft agar CTT plates (0.5% casitone, 10mM Tris-HCl pH 8.0, 1mM KPO_4_ pH 7.6, 8mM MgSO_4_, with the indicated concentrations of select agar (Invitrogen)) and incubated at 32°C for 24 hrs. Colony edges were imaged using a Leica MZ75 stereomicroscope with a Leica MC120 HD camera.

### Antibodies and immunoblot analysis

Immunoblotting was done with rabbit, polyclonal α-PilA and α-LonD antibodies^11^. As secondary antibodies goat, anti-rabbit immunoglobulin G peroxidase conjugate (Sigma-Aldrich, A8275) was used. Blots were developed using Luminata™ Western HRP substrate (Millipore).

### Data availability

The authors declare that all data supporting this study are available within the article, the Supplementary Information file or are available from the corresponding authors upon request. The source data underlying Fig. 1A, B; 2F; 5D, E, F; S2E, H; S4A; S5 are provided as a Source Data file.

## Acknowledgements

We thank Lisa Craig for providing grids with negatively-stained T4P from Ng and PAK.

## Funding

This work was supported by the National Fund for Scientific Research (FNRS) (to YFD), the Max Planck Society (to LSA), and NIH grant GM122510 (to EHE).

## Authors’ contributions

ATL: Designed and conceived the study, analyzed the length and phylogenetic distribution of T4 pilins (K02650), purified T4P, constructed and characterized *M. xanthus* strains expressing PilA^Mx^ variants, and determined PL for T4P from PilA^Mx^ variants. WZ: Performed the cryo-EM data collection, image processing and reconstruction, model building and refinement, and determined PL for T4P from Mx, Ng and Pa. AV: Performed the AFM force spectroscopy analyses and analyzed the data. SL: Generated plasmids and strains and performed motility assays. MH: Generated the AlphaFold-Multimer models and helped with the KEGG group sequence aquisition. YFD: Supervised AFM force spectroscopy and provided funding. LSA: Conceived the study, supervised research and provided funding. EHE: Conceived the study, supervised research and provided funding. ATL, LSA and EHE: Analyzed and interpreted data and wrote the manuscript. All authors approved the final manuscript.

## Competing interests

The authors declare no competing interests.

## Notes

### Competing Interest Statement

The authors have declared no competing interest.

## References

1 Wadhwa, N. & Berg, H. C. Bacterial motility: machinery and mechanisms. Nat Rev Microbiol 20, 161–173, (2022).

2 Miyata, M. et al. Tree of motility – A proposed history of motility systems in the tree of life. Genes Cells 25, 6–21, (2020).

3 Craig, L., Forest, K. T. & Maier, B. Type IV pili: dynamics, biophysics and functional consequences. Nat. Rev. Microbiol. 17, 429–440, (2019).

4 Giltner, C. L., Nguyen, Y. & Burrows, L. L. Type IV pilin proteins: versatile molecular modules. Microbiol. Mol. Biol. Rev. 76, 740–772, (2012).

5 Merz, A. J., So, M. & Sheetz, M. P. Pilus retraction powers bacterial twitching motility. Nature 407, 98–102, (2000).

6 Skerker, J. M. & Berg, H. C. Direct observation of extension and retraction of type IV pili. Proc Natl Acad Sci U S A 98, 6901–6904, (2001).

7 Clausen, M., Jakovljevic, V., Søgaard-Andersen, L. & Maier, B. High-force generation is a conserved property of type IV pilus systems. J Bacteriol 191, 4633–4638, (2009).

8 Maier, B., Potter, L., So, M., Seifert, H. S. & Sheetz, M. P. Single pilus motor forces exceed 100 pN. Proc Natl Acad Sci U S A 99, 16012–16017, (2002).

9 Gold, V. A., Salzer, R., Averhoff, B. & Kuhlbrandt, W. Structure of a type IV pilus machinery in the open and closed state. elife 4, e07380, (2015).

10 Chang, Y. W. et al. Architecture of the type IVa pilus machine. Science 351, aad2001, (2016).

11 Treuner-Lange, A. et al. PilY1 and minor pilins form a complex priming the type IVa pilus in *Myxococcus xanthus*. Nat Commun 11, 5054, (2020).

12 Bischof, L. F., Friedrich, C., Harms, A., Søgaard-Andersen, L. & van der Does, C. The type IV pilus assembly ATPase PilB of *Myxococcus xanthus* interacts with the inner membrane platform protein PilC and the nucleotide-binding protein PilM. J Biol Chem 291, 6946–6957, (2016).

13 Craig, L., Pique, M. E. & Tainer, J. A. Type IV pilus structure and bacterial pathogenicity. Nat Rev Microbiol 2, 363–378, (2004).

14 Herfurth, M. et al. A noncanonical cytochrome c stimulates calcium binding by PilY1 for type IVa pili formation. Proc Natl Acad Sci U S A 119, (2022).

15 Xue, S. et al. The differential expression of PilY1 proteins by the HsfBA phosphorelay allows twitching motility in the absence of exopolysaccharides. PLoS Genet 18, e1010188, (2022).

16 Rudel, T., Scheurerpflug, I. & Meyer, T. F. *Neisseria* PilC protein identified as type-4 pilus tip-located adhesin. Nature 373, 357–359, (1995).

17 Strom, M. S., Nunn, D. N. & Lory, S. A single bifunctional enzyme, PilD, catalyzes cleavage and N-methylation of proteins belonging to the type IV pilin family. Proc Natl Acad Sci U S A 90, 2404–2408, (1993).

18 Sheppard, D. et al. The major subunit of widespread competence pili exhibits a novel and conserved type IV pilin fold. J Biol Chem 295, 6594–6604, (2020).

19 Gu, Y. et al. Structure of *Geobacter* pili reveals secretory rather than nanowire behaviour. Nature 597, 430–434, (2021).

20 Wang, F., Craig, L., Liu, X., Rensing, C. & Egelman, E. H. Microbial nanowires: type IV pili or cytochrome filaments? Trends Microbiol 31, 384–392, (2023).

21 Wang, F. et al. Cryoelectron microscopy reconstructions of the *Pseudomonas aeruginosa* and *Neisseria gonorrhoeae* type IV pili at sub-nanometer resolution. Structure 25, 1423–1435 e1424, (2017).

22 Kolappan, S. et al. Structure of the *Neisseria meningitidis* type IV pilus. Nat Commun 7, 13015, (2016).

23 Bardiaux, B. et al. Structure and assembly of the enterohemorrhagic *Escherichia coli* type 4 pilus. Structure 27, 1082–1093 e1085, (2019).

24. Wang, F., et al. Cryo-EM structure of an extracellular Geobacter OmcE cytochrome filament reveals tetrahaem packing. Nat Microbiol 7, 1291–1300, (2022).

25 Lopez-Castilla, A. et al. Structure of the calcium-dependent type 2 secretion pseudopilus. Nat Microbiol 2, 1686–1695, (2017).

26 Neuhaus, A. et al. Cryo-electron microscopy reveals two distinct type IV pili assembled by the same bacterium. Nat Commun 11, 2231, (2020).

27 Craig, L. et al. Type IV pilus structure by cryo-electron microscopy and crystallography: implications for pilus assembly and functions. Mol Cell 23, 651–662, (2006).

28 Touhami, A., Jericho, M. H., Boyd, J. M. & Beveridge, T. J. Nanoscale characterization and determination of adhesion forces of *Pseudomonas aeruginosa* pili by using atomic force microscopy. J Bacteriol 188, 370–377, (2006).

29 Biais, N., Higashi, D. L., Brujic, J., So, M. & Sheetz, M. P. Force-dependent polymorphism in type IV pili reveals hidden epitopes. Proc Natl Acad Sci U S A 107, 11358–11363, (2010).

30 Beaussart, A. et al. Nanoscale adhesion forces of *Pseudomonas aeruginosa* type IV pili. ACS Nano 8, 10723–10733, (2014).

31 Brissac, T., Mikaty, G., Duménil, G., Coureuil, M. & Nassif, X. The meningococcal minor pilin PilX is responsible for type IV pilus conformational changes associated with signaling to endothelial cells. Infect Immun 80, 3297–3306, (2012).

32 Wu, S. S. & Kaiser, D. Genetic and functional evidence that Type IV pili are required for social gliding motility in *Myxococcus xanthus*. Mol Microbiol 18, 547–558, (1995).

33 Bader, C. D., Panter, F. & Müller, R. In depth natural product discovery – Myxobacterial strains that provided multiple secondary metabolites. Biotechnol Adv. 39, 107480, (2020).

34 Konovalova, A., Petters, T. & Søgaard-Andersen, L. Extracellular biology of *Myxococcus xanthus*. FEMS Microbiol. Rev. 34, 89–106, (2010).

35 Zhang, Y., Ducret, A., Shaevitz, J. & Mignot, T. From individual cell motility to collective behaviors: insights from a prokaryote, *Myxococcus xanthus*. FEMS Microbiol. Rev. 36, 149–164, (2012).

36 Shi, W. & Zusman, D. R. The two motility systems of *Myxococcus xanthus* show different selective advantages on various surfaces. Proc Natl Acad Sci U S A 90, 3378–3382, (1993).

37 Kanehisa, M., Sato, Y., Kawashima, M., Furumichi, M. & Tanabe, M. KEGG as a reference resource for gene and protein annotation. Nucleic Acids Res 44, D457–462, (2016).

38 Garrity, G. M., Bell, J. A. & Lilburn, T. in Bergey’s Manual of Systematics of Archaea and Bacteria (2015).

39 Coenye, T. & Vandamme, P. Diversity and significance of *Burkholderia* species occupying diverse ecological niches. Environ Microbiol 5, 719–729, (2003).

40 Martinucci, M. et al. Accurate identification of members of the *Burkholderia cepacia* complex in cystic fibrosis sputum. Lett Appl Microbiol 62, 221–229, (2016).

41 Ligthart, K., Belzer, C., de Vos, W. M. & Tytgat, H. L. P. Bridging bacteria and the gut: Functional aspects of type IV pili. Trends Microbiol 28, 340–348, (2020).

42 Rozman, V., Accetto, T., Duncan, S. H., Flint, H. J. & Vodovnik, M. Type IV pili are widespread among non-pathogenic Gram-positive gut bacteria with diverse carbohydrate utilization patterns. Environ Microbiol 23, 1527–1540, (2021).

43 Orans, J. et al. Crystal structure analysis reveals *Pseudomonas* PilY1 as an essential calcium-dependent regulator of bacterial surface motility. Proc Natl Acad Sci U S A 107, 1065–1070, (2010).

44 Remaut, H. & Waksman, G. Protein-protein interaction through beta-strand addition. Trends Biochem Sci 31, 436–444, (2006).

45 Vetsch, M. et al. Mechanism of fibre assembly through the chaperone-usher pathway. EMBO Rep 7, 734–738, (2006).

46 Zyla, D., Echeverria, B. & Glockshuber, R. Donor strand sequence, rather than donor strand orientation, determines the stability and non-equilibrium folding of the type 1 pilus subunit FimA. J Biol Chem 295, 12437–12448, (2020).

47 Galkin, V. E., Orlova, A., Vos, M. R., Schroder, G. F. & Egelman, E. H. Near-atomic resolution for one state of f-actin. Structure 23, 173–182, (2015).

48 Galkin, V. E., Orlova, A. & Egelman, E. H. Actin filaments as tension sensors. Current Biology 22, R96–R101, (2012).

49 Lu, S. et al. Nanoscale pulling of type IV pili reveals their flexibility and adhesion to surfaces over extended lengths of the pili. Biophys J 108, 2865–2875, (2015).

50 Kaiser, D. Social gliding is correlated with the presence of pili in *Myxococcus xanthus*. Proc. Natl. Acad. Sci. USA 76, 5952–5956, (1979).

51 Cont, A., Vermeil, J. & Persat, A. Material substrate physical properties control *Pseudomonas aeruginosa* biofilm architecture. mBio, e0351822, (2023).

52 Yang, Z. et al. Alanine 32 in PilA is important for PilA stability and type IV pili function in *Myxococcus xanthus*. Microbiology (Reading) 157, 1920–1928, (2011).

53 Yang, Z., Lux, R., Hu, W., Hu, C. & Shi, W. PilA localization affects extracellular polysaccharide production and fruiting body formation in *Myxococcus xanthus*. Mol Microbiol 76, 1500–1513, (2010).

54 Nivaskumar, M. et al. Distinct docking and stabilization steps of the Pseudopilus conformational transition path suggest rotational assembly of type IV pilus-like fibers. Structure 22, 685–696, (2014).

55 Karami, Y. et al. Computational and biochemical analysis of type IV pilus dynamics and stability. Structure 29, 1397–1409 e1396, (2021).

56 Campos, M., Nilges, M., Cisneros, D. A. & Francetic, O. Detailed structural and assembly model of the type II secretion pilus from sparse data. Proc Natl Acad Sci U S A 107, 13081–13086, (2010).

57 Koch, M. D., Fei, C., Wingreen, N. S., Shaevitz, J. W. & Gitai, Z. Competitive binding of independent extension and retraction motors explains the quantitative dynamics of type IV pili. Proc Natl Acad Sci U S A 118, (2021).

58 Tala, L., Fineberg, A., Kukura, P. & Persat, A. *Pseudomonas aeruginosa* orchestrates twitching motility by sequential control of type IV pili movements. Nat Microbiol 4, 774–780, (2019).

59 Aoki, K. F. & Kanehisa, M. Using the KEGG database resource. Curr. Protoc. Bioinformatics 11, 1.12.11–11.12.54, (2005).

60 Li, W. & Godzik, A. Cd-hit: a fast program for clustering and comparing large sets of protein or nucleotide sequences. Bioinformatics 22, 1658–1659, (2006).

61 Teufel, F. et al. SignalP 6.0 predicts all five types of signal peptides using protein language models. Nat Biotechnol 40, 1023–1025, (2022).

62 Pei, J., Tang, M. & Grishin, N. V. PROMALS3D web server for accurate multiple protein sequence and structure alignments. Nucleic Acids Res 36, W30–34, (2008).

63 Notredame, C., Higgins, D. G. & Heringa, J. T-Coffee: A novel method for fast and accurate multiple sequence alignment. J. Mol. Biol. 302, 205–217, (2000).

64 Thompson, J. D., Higgins, D. G. & Gibson, T. J. CLUSTAL W: improving the sensitivity of progressive multiple sequence alignment through sequence weighting, position-specific gap penalties and weight matrix choice. Nucleic Acids Res. 22, 4673–4680, (1994).

65. Hall, T. A. BioEdit: A user-friendly biological sequence alignment editor and analysis program for Windows 95/98/NT. Nucleic Acids Symposium Series 41, 95–98, (1999).

66 Cai, W., Pei, J. & Grishin, N. V. Reconstruction of ancestral protein sequences and its applications. BMC Evol Biol 4, 33, (2004).

67 Gabler, F. et al. Protein sequence analysis using the MPI Bioinformatics Toolkit. Curr Protoc Bioinformatics 72, e108, (2020).

68 Letunic, I. & Bork, P. Interactive Tree Of Life (iTOL): an online tool for phylogenetic tree display and annotation. Bioinformatics 23, 127–128, (2007).

69 Shi, X. et al. Bioinformatics and experimental analysis of proteins of two-component systems in *Myxococcus xanthus*. J. Bacteriol. 190, 613–624, (2008).

70 Søgaard-Andersen, L., Slack, F. J., Kimsey, H. & Kaiser, D. Intercellular C-signaling in *Myxococcus xanthus* involves a branched signal transduction pathway. Genes Dev. 10, 740–754, (1996).

71. 71 Sambrook, J. & Russell, D. W. Molecular cloning : a laboratory manual. 3rd edn, (Cold Spring Harbor Laboratory Press, 2001).

72 Wu, S. S., Wu, J. & Kaiser, D. The *Myxococcus xanthus pilT* locus is required for social gliding motility although pili are still produced. Mol. Microbiol. 23, 109–121, (1997).

73 Pettersen, E. F. et al. UCSF Chimera--a visualization system for exploratory research and analysis. J Comput Chem 25, 1605–1612, (2004).

74 Emsley, P., Lohkamp, B., Scott, W. G. & Cowtan, K. Features and development of Coot. Acta Crystallogr D Biol Crystallogr 66, 486–501, (2010).

75 Adams, P. D. et al. PHENIX: a comprehensive Python-based system for macromolecular structure solution. Acta Crystallogr D Biol Crystallogr 66, 213–221, (2010).

76 Chen, V. B. et al. MolProbity: all-atom structure validation for macromolecular crystallography. Acta Crystallogr D Biol Crystallogr 66, 12–21, (2010).

77. Evans, R., et al. Protein complex prediction with AlphaFold-Multimer. *bioRxiv*, 10.1101/2021.1110.1104.463034., (2022).

78 Jumper, J. et al. Highly accurate protein structure prediction with AlphaFold. Nature 596, 583–589, (2021).

79 Mirdita, M. et al. ColabFold: making protein folding accessible to all. Nat Methods 19, 679–682, (2022).

80 Schwabe, J., Pérez-Burgos, M., Herfurth, M., Glatter, T. & Søgaard-Andersen, L. Evidence for a widespread third system for bacterial polysaccharide export across the outer membrane comprising a composite OPX/beta-barrel translocon. mBio 13, e0203222, (2022).

81 Krissinel, E. & Henrick, K. Inference of macromolecular assemblies from crystalline state. J Mol Biol 372, 774–797, (2007).

82 Collins, T. J. ImageJ for microscopy. Biotechniques 43, 25–30, (2007).

83 Hutter, J. L. & Bechhoefer, J. Calibration of atomic-force microscope tips. Review of Scientific Instruments 64, 1868–1873, (1993).

