## supplemental files for "Large pilin subunits provide distinct structural and mechanical properties for the *Myxococcus xanthus* type IV pilus"

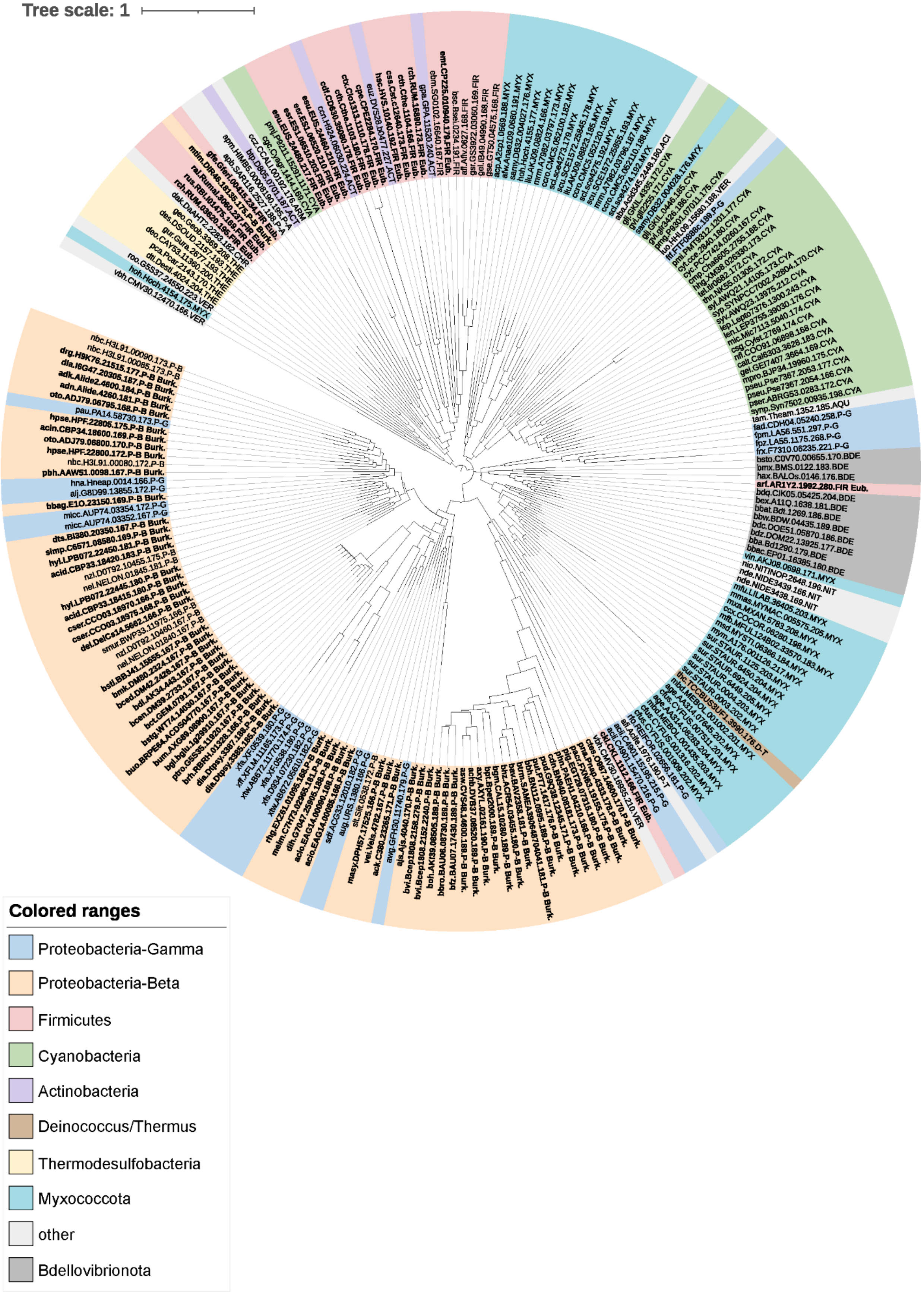
Tree scale: 1
Colored ranges
Proteobacteria-Gamma
Proteobacteria-Beta
Firmicutes
Cyanobacteria
Actinobacteria
Deinococcus/Thermus
Thermodesulfobacteria
Myxococcota
other
Bdellovibrionota

**Figure S1. Phylogenetic tree of 226 large major pilins.**

Phylogenetic tree of 226 large major pilins, as shown in Table S1 and listed in Table S2. The color code (bottom left corner) is the same as used in Table S1. The large pilins of the Betaproteobacteria are found especially in Burkholderiales, and those are shown in bold with the abbreviation **Burk**. at the end of the locus tag.

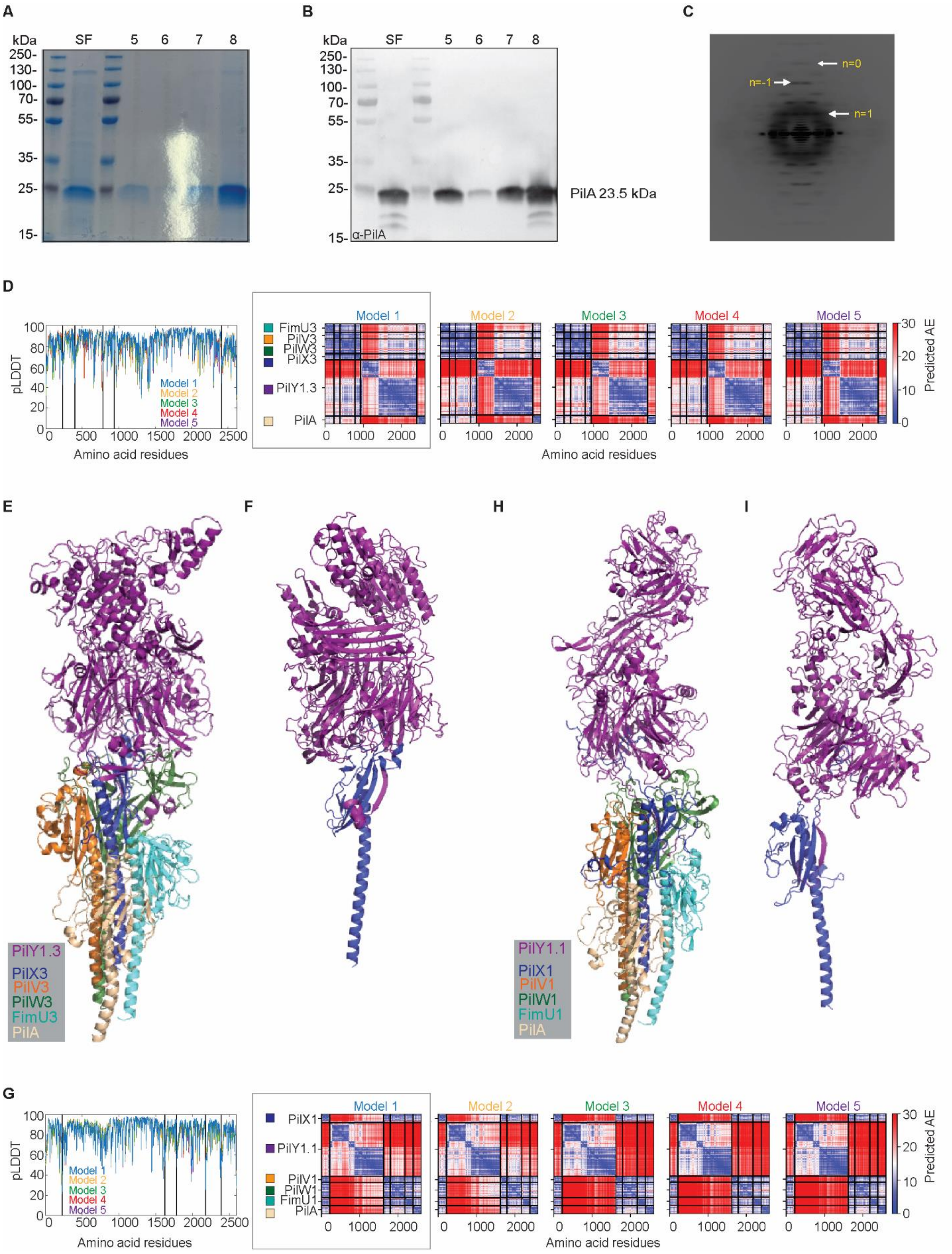
A
kDa
SF
5
6
7
8
250-
130-
100-
70-
55-
35-
25-
15-
B
α-PilA
PilA 23.5 kDa
C
n=0
n=-1
n=1
D
pLDDT
Amino acid residues
Model 1
Model 2
Model 3
Model 4
Model 5
FimU3
PilV3
PilW3
PilX3
PilY1.3
PilA
Predicted AE
E
F
H
I
PilY1.3
PilX3
PilV3
PilW3
FimU3
PilA
PilY1.1
PilX1
PilV1
PilW1
FimU1
PilA
G
PilX1
PilY1.1
PilV1
PilW1
FimU1
PilA

**Figure S2. Structure elucidation of T4P$^{Mx}$ and AlphaFold modeling of two tip complexes.**
**A-B.** T4P$^{Mx}$ from Δ*pilT* cells were sheared-off and precipitated (sheared fraction = SF).
The SF was further purified using sucrose-gradient centrifugation and fractions were collected. SF and PilA-containing fractions were separated by SDS-PAGE and visualized by Coomassie protein staining (**A**) and probed with α-PilA antibodies (**B**). The calculated molecular mass of PilA and positions of molecular markers are indicated.
**C.** Averaged power spectrum from T4P$^{Mx}$.
**D**. Plots generated by AlphaFold for the model of cluster_3 proteins (FimU3, PilV3, PilW3, PilV3, and PilY1.3) and PilA with pLDDT (left) and pAE plots (right) for five models of the indicated proteins as predicted by AlphaFold. Model 1-5 in the pLDDT (left) and pAE plots (right) are shown in the same colors. Model 1 (marked by a grey box) was used for analysis.
**E**. AlphaFold model of cluster_3 proteins (as in **D**) and PilA in the indicated colors.
**F**. Predicted protein-protein interaction by β-strand addition between PilY1.3 (purple) and PilX3 (blue) by the AlphaFold model of cluster_3 proteins.
**G**. Plots generated by AlphaFold for the model of cluster_1 proteins (FimU1, PilV1, PilW1, PilX1, and PilY1.1) and PilA with pLDDT (left) and pAE plots (right) for five models of the indicated proteins as predicted by AlphaFold. Model 1-5 in the pLDDT (left) and pAE plots (right) are shown in the same colors. Model 1 (marked by a grey box) was used for analysis.
**H**. AlphaFold model of cluster_1 proteins (as in **G**) and PilA in the indicated colors.
**I.** Predicted protein-protein interaction by β-strand addition between PilY1.1 (purple) and PilX1 (blue) by the AlphaFold model of cluster_1 proteins.
**D-I.** The sequences of the mature minor pilins, and PilY1 proteins without their signal peptides as reported earlier[1], were used for generating the models.

**A**

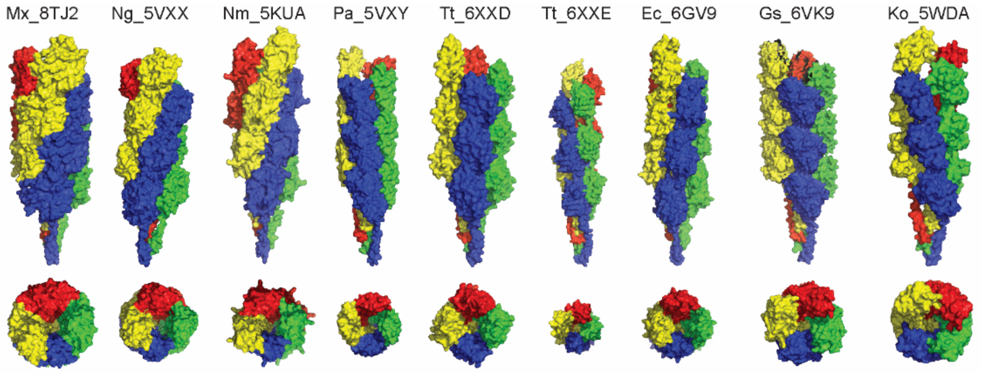


| | Mx_8TJ2 | Ng_5VXX | Nm_5KUA | Pa_5VXY | Tt_6XXD | Tt_6XXE | Ec_6GV9 | Gs_6VK9 | Ko_5WDA |
|---|---|---|---|---|---|---|---|---|---|
| resolution (Å): | 3.0 | 5.1 | 6.0 | 8.0 | 3.2 | 3.5 | 8.0 | 3.8 | 5.0 |
| diameter (Å): | ~70 | ~62 | ~60 | ~52 | 70 | 45 | ~60 | ~65 | ~70 |
| rise (Å): | 10 | 10.1 | 10.3 | 10.5 | 9.3 | 11.3 | 11.2 | 10.4 | 10.2 |
| twist (°): | 100.7 | 100.8 | 100.8 | 87.6 | 92.5 | 84.3 | 96 | 89.1 | 83.5 |
| subunits / turn: | 3.6 | 3.6 | 3.6 | 4.1 | 3.9 | 4.3 | 3.8 | 4.0 | 4.3 |
| pilin length (aa): | 208 | 158 | 161 | 144 | 125 | 111 | 140 | 165 | 134 |
| persistence length (µm): | 21 | 11 | - | 13 | - | - | - | - | - |

**B**

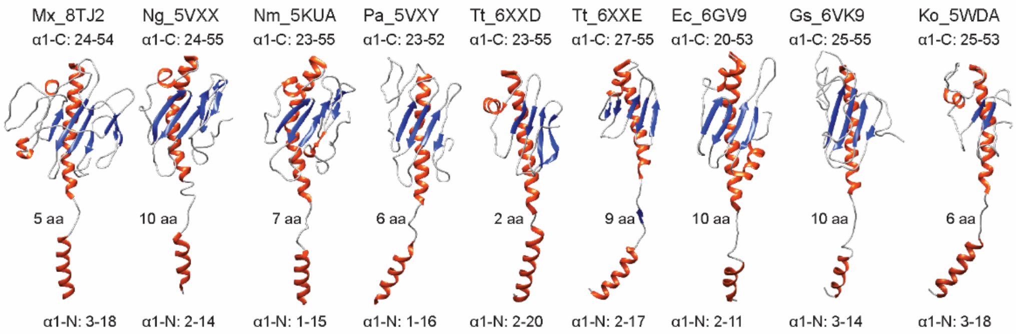


| | Mx_8TJ2 | Ng_5VXX | Nm_5KUA | Pa_5VXY | Tt_6XXD | Tt_6XXE | Ec_6GV9 | Gs_6VK9 | Ko_5WDA |
|---|---|---|---|---|---|---|---|---|---|
| pilin length (aa): | 208 | 158 | 161 | 144 | 125 | 111 | 140 | 165 | 134 |
| % structured (α+β): | 41.3 | 51.3 | 71.7 | 42.4 | 65.5 | 62.2 | 57.1 | 38.3 | 43.3 |
| globular domain (aa): | 153 | 103 | 106 | 92 | 70 | 56 | 87 | 110 | 81 |
| % structured (α+β): | 25.5 | 35.9 | 31.1 | 17.6 | 42.8 | 41.5 | 41.4 | 19.1 | 14.8 |

**C**

```
                           α1-N                          α1-C                                α2
Mx PilA
Mx PilA     FTLIELMIVVAIIGILAAIAIPNFIKFQARSKQSEAKTNLKALYTAQKSFFSEKDRYSDF-----------AN----------E  63
Ng PilE     FTLIELMIVIAIVGILAAVALPAYQDYTARAQVSEAILLAEGQKSAVTEYYLNHGKWPEN-----------NT----------S  63
Nm PilE     FTLIELMIVIAIVGILAAVALPAYQDYTARAQVSEAILLAEGQKSAVTEYYLNHGEWPGD-----------NS----------S  63
Pa PilA      FTLIELMIVVAIIGILAAIAIPQYQNYVARSEGASALASVNPLKTTVEEALSRGWSVKSGTGTEDATKKEVPLGVAADANKLGT  84
Tt PilA4    FTLIELLIVIAIIAILAAVLIPNLLAARKRANDTVVTAYLNDAVKFQEMYQIDNNSYTSN-----------QAALI-------S  66
Tt PilA5    FTLIELAIVIVIIGILVAIAVPRFVDLTDQANQANVDATAAAVRSAYAIATVQAKGIPTC-----------DQ-----------  62
Ec PpdD     FTLIELMVVIGIIAILSAIGIPAYQNYLRKAALTDMLQTFVPYRTAVELCALEHGGLDTC-----------DGG---------S  64
Gs PilA-N/C FTLIELLIVVAIIGILAAIAIPQFSAYRVKAYNSAASSDLRNLKTALESAFADDQTYPPE-----------SAGKIPTTTMGGK  73
Ko PulG     FTLLEIMVVIVILGVLASLVVPNLMGNKEKADRQKVVSDLVALEGALDMYKLDNSRYPTT-----------EQGL--------Q  65

                  β1                                 α3   α4                 β1'
Mx PilA
Mx PilA     IGFAPERG---NRYGYRVSAAAGDCEVRNAADLPVPAAGVPCISNDSFRFGANS--AIDDP-----TPVVARFVPQGAAGWNTT 137
Ng PilE     AGVASPPSDIKGKYVKEVE----------------VKNGVVT--ATMLSSGVNN--EIKGK-----KLSLWARRENGSVKWFCG 122
Nm PilE     AGVAT-SADIKGKYVQSVT----------------VANGVIT--AQMASSNVNN--EIKSK-----KLSLWAKRQNGSVKWFCG 121
Pa PilA     I-------------ALKPD----------------PADGTADITLTFTMGGAGP-------KNKGKIITLT-RT-AADGLWKCT 130
Tt PilA4    LGLKSTPAN---VTF-----------------------SIVS--AS-------------AN-----SYCMI-AGHSGGTVWFAA 103
Tt PilA5    -----------------------------------------------VF--ANP--EGGST-----SGSTWTSSDNS-TTVSCN  89
Ec PpdD     NGIPSPTT---TRYVSAMS----------------VAKGVVS--LTGQE-SLN---GLSVV-----MTPGW-DNANGVTGWARN 117
Gs PilA-N/C DFTFKPSTN---VSVSYF-----------------TTNGATSTAG-----TVNTDYAVNTKNSSGN-RVFT-STNNTSNIWYIE 130
Ko PulG     ALVSAPSAE---PHARNY-----------------PEGGYIR--RLP------Q--DPWGS-----DYQLLSPGQCGQVDIFSL 114

                      β2           β3      β2'       β3'         β4
Mx PilA
Mx PilA     LGVQ-PTIADCPNCNFFAGARGNADNEATFDDWVIAGFEGSGQVGPCSEAGNVASG--TP--YNTR----NDVACD--G-AAQ 208
Ng PilE     QPVT-RTD------DD-TVADAKDGKEID------------------------TKH--LP--STCR----DNFDA---K---- 158
Nm PilE     QPVT-RTTAT---ATDVAAANGKTDDKIN------------------------TKH--LP--STCR----DDSSA---S---- 161
Pa PilA     -------------------------SDQD------------------------EQF--IP--KGCS----------------R 144
Tt PilA4    TPDK-GVY--------------KTNTAVT------------------------S-S--QP--ESCP----------------- 125
Tt PilA5    A-------------------------SAD------------------------TFT--IS--RGGKTRTLNLTVN-------- 111
Ec PpdD     ------------------------CNIQS------------------------DSA--LQ--QACE----DVFRFDDAN---- 140
Gs PilA-N/C NDAWKGK------------------AVSD------------------------SDVTALGTGDVGK----SDFSGT--EWKSQ 165
Ko PulG     GPD--GVP--------------ESNDDIG------------------------NCT--IG--KK------------------- 134
```

**Figure S3. Comparisons of T4P$^{Mx}$ and PilA$^{Mx}$ to other T4P structures and T4 pilins.**

**A**. Helical arrangement of depicted T4P with N and N+4 subunits shown in the same color. T4P characteristics (resolution, diameter, rise, twist, subunits/turn and pilin length were taken from this study and previously solved T4P structures (PilE from *N. meningitidis* (Nm PilE; 5KUA)[2]; PilA from *P. aeruginosa* PAK (Pa PilA; 5VXY)[3]; PilE from *N. gonorrhoeae* (Ng PilE; 5VXX)[3]; PilA4 (Tt PilA4; 6XXD) and PilA5 (Tt PilA5; 6XXE) from *T. thermophilus*[4]; PilA-N from *G. sulfurreducens* (Gs PilA-N; 6VK9)[5]; PpdD from enterohemorrhagic *E. coli* (Ec PpdD; 6GV9)[6], and PulG from *K. oxytoca* (Ko PulG; 5WDA)[7]. The last row indicates persistence length as measured and depicted in Fig. S4A.

**B**. Ribbon representation of PilA$^{Mx}$ and depicted pilins from previously solved T4P structures as in **A** with helical elements (α) shown in red, β-stranded elements (β) shown in blue and less-structured areas (loops) in grey. The first and last residues of the helices of α1-N and α1-C are shown as well as the number of residues (aa) in the melted region. Pilin characteristics (length of pilin and globular domain (aa) and % structured (α+β) were taken from this study and previously solved T4P structures as in **A**.

**C** Multiple sequence alignment of PilA$^{Mx}$ and pilins from previously solved T4P structures as in **A**. The top row indicates the structural elements of PilA$^{Mx}$ as in Fig. 3B.

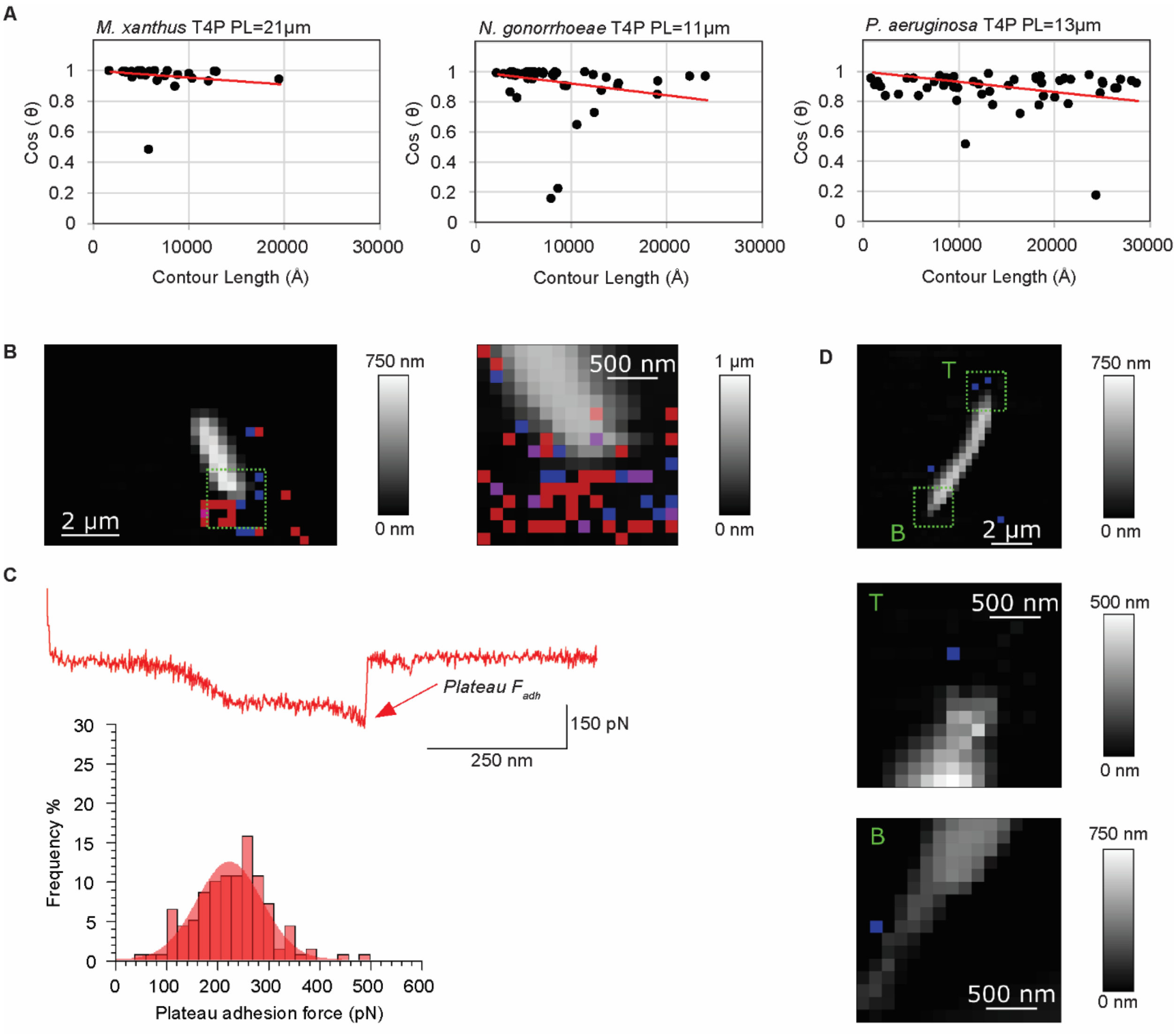


**Fig. S4. Persistence length analyses of T4P and AFM-MS analyses of *M. xanthus* cells.**
**A** The cosine of the bending angle (theta) of a T4P segment is plotted against the contour length of the segment (Å). PL (µm) is calculated from the slope of the linear trendline. Negative staining micrographs of purified T4P from *M. xanthus* (left, as in Fig. 2A), T4P from *N. gonorrhoeae* (center), and T4P from *P. aeruginosa* PAK (right) were used for this analysis.
**B**. *M. xanthus* WT cells probed in force volume mode in which an *F-d* curve is recorded at each pixel of a rectangular raster grid using a constant approach and retract velocity. Shown is a representative height map of a whole cell (left) and zoom of the lower pole (right, 16 × 16 pixels). The overlayed red, blue, and purple pixels indicate where force plateau, nanospring or both signatures were detected.

**C.** Representative *F-d* curve obtained for a *M. xanthus* WT cell showing a force plateau signature (top). The rupture/adhesion force ($F_{adh}$) is indicated by a red arrow. Bottom: Histogram showing the distribution of plateau $F_{adh}$ as determined from *F-d* curves generated at a probe retraction velocity of 5µm/sec (n= 141 force plateaus in 125/5400 curves from 5 tip-cell combinations). Number indicates mean ± STDEV.

**D.** Overlayed adhesion and height images of *M. xanthus* ∆*pilA* cells obtained using the same force spectroscopy parameters used for WT in panel **B**. Top: Representative cell, middle: zoom of the pole at the top (T) (16 × 16 pixels), bottom: zoom of the pole at the bottom (B) (bottom, 16 × 16 pixels). Signatures resembling nanosprings were detected sporadically. Plateau signatures were not detected.

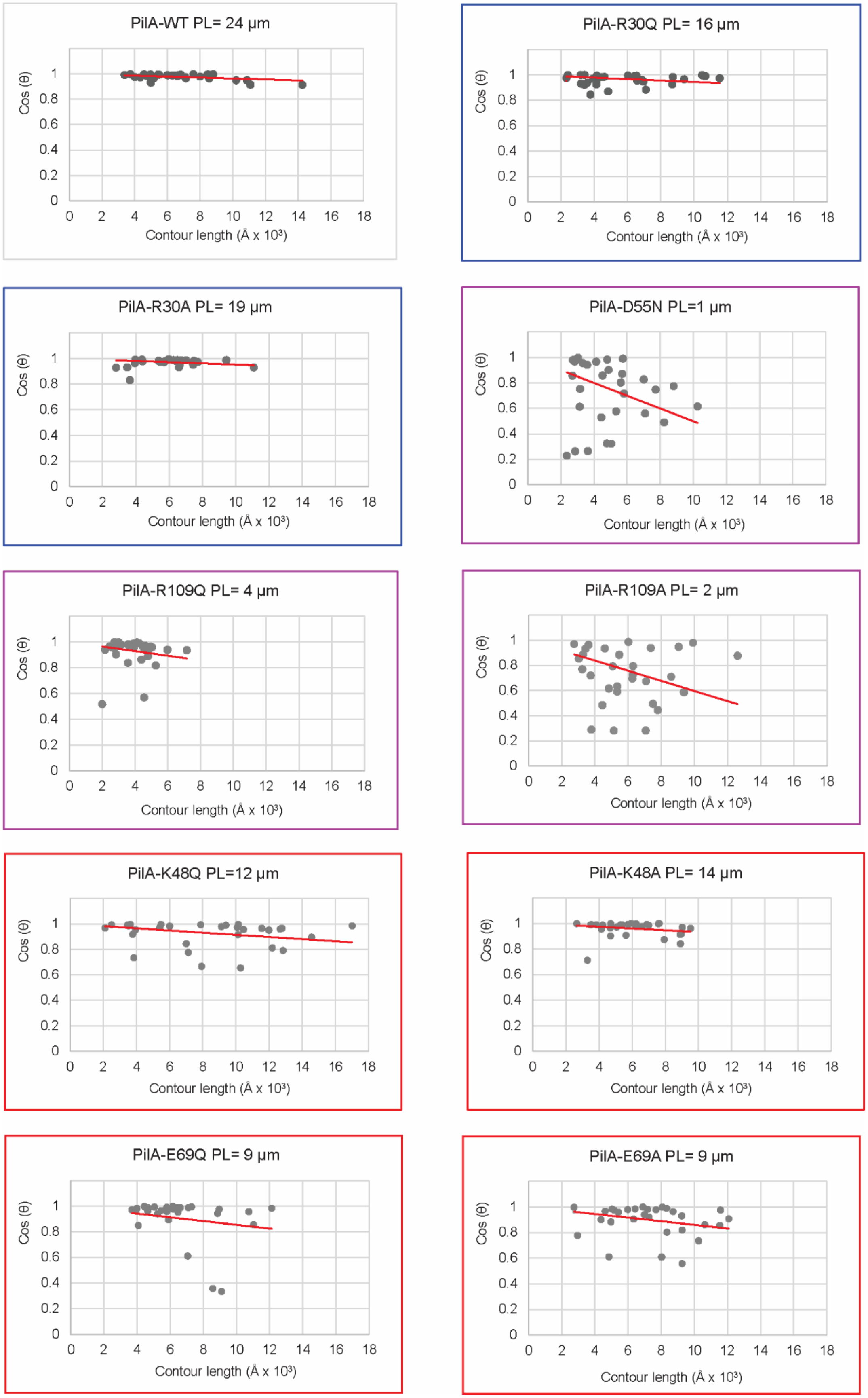
PilA-WT PL= 24 µm
PilA-R30Q PL= 16 µm
PilA-R30A PL= 19 µm
PilA-D55N PL=1 µm
PilA-R109Q PL= 4 µm
PilA-R109A PL= 2 µm
PilA-K48Q PL=12 µm
PilA-K48A PL= 14 µm
PilA-E69Q PL= 9 µm
PilA-E69A PL= 9 µm
Cos (θ)
Contour length (Å x 10³)
0
0.2
0.4
0.6
0.8
1
0 2 4 6 8 10 12 14 16 18

**Figure S5. Persistence length analyses.**

The cosine of the bending angle (theta) of a T4P segment is plotted against the contour length of the segment (Å). PL (µm) is calculated from the slope of the linear trendline. Negative staining micrographs of sheared pili from ∆*pilT* strains expressing $PilA^{WT}$ and indicated PilA variants as shown in Fig. 5F were used for this analysis.

**Table S1. Phylogenetic distribution of major pilins and the large major pilins subset.**

| | phyla (proteobacteria shown as class) | n =1955 | % of 1955 | ≥166 aa | % of 226 |
|---|---|---|---|---|---|
| P-G | Proteobacteria-Gamma | 707 | 36.2 | 22 | 9.7 |
| P-B | Proteobacteria-Beta | 374 | 19.1 | 75 | 33.2 |
| FIR | Firmicutes | 339 | 17.3 | 25 | 11.1 |
| CYA | Cyanobacteria | 126 | 6.4 | 29 | 12.8 |
| ACT | Actinobacteria | 72 | 3.7 | 4 | 1.8 |
| D-T | Deinococcus-Thermus | 70 | 3.6 | 1 | 0.4 |
| THE | Thermodesulfobacteria | 56 | 2.9 | 7 | 3.1 |
| MYX | Myxococcota | 42 | 2.1 | 39 | 17.3 |
| TTG | Thermotogae | 23 | 1.2 | | 0.0 |
| | other (each less than 2%) | 146 | 7.5 | 24 | 10.6 |
| | | **1955** | | **226** | |
| VER | Verrumicrobia | 17 | 0.9 | 4 | 1.8 |
| AQU | Aquificae | 17 | 0.9 | 1 | 0.4 |
| ACI | Acidobacteria | 16 | 0.8 | 1 | 0.4 |
| CHL | Chloroflexi | 15 | 0.8 | | 0.0 |
| FUS | Fusobacteria | 14 | 0.7 | | 0.0 |
| P-T | Proteobacteria-Acidithiobacilli | 12 | 0.6 | 1 | 0.4 |
| BDE | Bdellovibrionota | 11 | 0.6 | 11 | 4.9 |
| PLA | Planctomycetes | 9 | 0.5 | | 0.0 |
| CAM | Campylobacterota | 9 | 0.5 | | 0.0 |
| P-A | Proteobacteria-Alpha | 6 | 0.3 | 2 | 0.9 |
| NIT | Nitrospirae | 6 | 0.3 | 3 | 1.3 |
| GEM | Gemmatimonadetes | 5 | 0.3 | | 0.0 |
| DES | Desferribacteres | 4 | 0.2 | | 0.0 |
| ARM | Armatimonadetes | 3 | 0.2 | 1 | 0.4 |
| TEN | Tenericutes | 2 | 0.1 | | 0.0 |
| | | **146** | | **24** | |

The dataset (n=1955) contains all mature pilin sequences of the KEGG group K02650 (type IV pilus assembly protein PilA), after filtering out highly homologous sequences (> 90% identity), sequences lacking a T3SP and a classified taxonomy. Bacterial phyla, with sequences derived from only one genome, were also excluded from the analysis. The first row indicates the color code and the abbreviations of phyla or proteobacterial classes listed in the 2$^{nd}$ row. The grey shaded field in the top part indicates the number of sequences in the category called "other". The number of sequences from these phyla and proteobacterial classes is shown in detail in the bottom part of the table and represents each less than 2 % of the 1,955 sequences. Please note that in Fig. 1A this category is presented in two columns called "other" and Bdellovibrionota. The shaded fields (last row) indicate phyla or proteobacterial classes, where the abundance of the 226 large major pilins (last row, % of 226) is overrepresented in comparison to their abundance in the complete dataset (4th row, % of 1955).

**Table S2. Phylogenetic distribution of large major pilins in the data subset.**

| phyla (proteobacteria as classes) | no. | Order | *Species* | locus tag | ength* (aa) |
|---|---|---|---|---|---|
| Proteobacteria-Gamma | 22 | Acidoferrobacterales | *Acidiferrobacter sp. SPIII_3* | acii_C4901_15195 | 215 |
| | | | | acii_C4901_15470 | 216 |
| | | Moraxellales | *Acinetobacter lanii* | alj_G8D99_13855 | 172 |
| | | | *Acinetobacter ursingii* | aug_URS_1380 | 166 |
| | | | *Acinetobacter wanghuae* | awg_GFH30_11740 | 179 |
| | | Thiotrichales | *Francisella adeliensis* | fad_CDH04_05240 | 258 |
| | | | *Francisella philomiragia O#319-036* | fpm_LA56_551 | 297 |
| | | | *Francisella philomiragia GA01-2794* | fpz_LA55_1175 | 268 |
| | | | *Francisella uliginis* | frx_F7310_06235 | 221 |
| | | | *Francisella tularensis subsp. tularensis FSC198* | ftf_FTF0888c | 189 |
| | | Chromatiales | *Halothiobacillus neapolitanus* | hna_Hneap_0014 | 166 |
| | | Cellvibrionales | *Microbulbifer aggregans* | micc_AUP74_03352 | 167 |
| | | | | micc_AUP74_03354 | 172 |
| | | Pseudomonadales | *Pseudomonas aeruginosa UCBPP-PA14* | pau_PA14_58730 | 173 |
| | | Oceanopspirillales | *Reinekea forsetii* | rfo_REIFOR_02656 | 181 |
| | | Nevskiales | *Steroidobacter denitrificans* | sdf_ACG33_12010 | 182 |
| | | Xanthomonadales | *Xylella fastidiosa 9a5c* | xfa_XF_0538_ 180 | 180 |
| | | | | xfa_XF_0539_ 180 | 180 |
| | | | *Xylella fastidiosa subsp. fastidiosa GB514* | xff_XFLM_11165 | 173 |
| | | | *Xylella fastidiosa subsp. sandyi Ann-1* | xfs_D934_07230 | 182 |
| | | | *Xylella taiwanensis* | xtw_AB672_05610 | 182 |
| | | | | xtw_AB672_11770 | 174 |
| Proteobacteria-Beta | 75 | Burkholderiales | *Achromobacter sp. B7* | achb_DVB37_08520 | 189 |
| | | | *Achromobacter spanius* | asw_CVS48_14600 | 189 |
| | | | *Achromobacter xylosoxidans A8* | axy_AXYL_02161 | 190 |
| | | | *Acidovorax carolinensis NA2* | acid_CBP33_18415 | 180 |
| | | | | acid_CBP33_18420 | 183 |
| | | | *Acidovorax carolinensis NA3* | acin_CBP34_18600 | 169 |
| | | | *Acidovorax ebreus* | dia_Dtpsy_3385 | 185 |
| | | | | dia_Dtpsy_3387 | 188 |
| | | | *Acidovorax sp. 1608163* | acio_EAG14_00085 | 166 |
| | | | | acio_EAG14_00090 | 168 |
| | | | *Acidovorax sp. JS42* | ajs_Ajs_4040 | 170 |
| | | | *Acidovorax sp. KKS102* | ack_C380_23265 | 171 |
| | | | *Alicycliphilus denitrificans BC* | adn_Alide_4260 | 181 |
| | | | *Alicycliphilus denitrificans K601* | adk_Alide2_4600 | 184 |
| | | | *Bordetella avium* | bav_BAV2358 | 181 |

| | | | | | |
|---|---|---|---|---|---|
| | | | *Bordetella bronchialis* | bbro_BAU06_08730 | 189 |
| | | | *Bordetella bronchiseptica 253* | bbh_BN112_0995 | 189 |
| | | | *Bordetella flabilis* | bfz_BAU07_17430 | 189 |
| | | | *Bordetella genomosp. 13* | bgm_CAL15_10280 | 189 |
| | | | *Bordetella hinzii* | bhz_ACR54_03455 | 180 |
| | | | *Bordetella petrii* | bpt_Bpet2000 | 189 |
| | | | *Bordetella sp. H567* | boh_AKI39_08505 | 189 |
| | | | *Bordetella trematum* | btrm_SAMEA390648 | 181 |
| | | | *Burkholderia cenocepacia DDS 22E-1* | bcen_DM39_2733 | 167 |
| | | | *Burkholderia cepacia DDS 7H-2* | bced_DM42_2426 | 167 |
| | | | *Burkholderia cepacia GG4* | bct_GEM_0791 | 167 |
| | | | *Burkholderia dolosa* | bdl_AK34_443 | 167 |
| | | | *Burkholderia glumae BGR1* | bgl_bglu_1g29910 | 167 |
| | | | *Burkholderia multivorans DDS 15A-1* | bmk_DM80_2324_ | 167 |
| | | | *Burkholderia sp. PAMC 26561* | bum_AXG89_08900 | 167 |
| | | | *Burkholderia stabilis* | bstl_BBJ41_15555 | 167 |
| | | | *Burkholderia stagnalis* | bstg_WT74_14030 | 167 |
| | | | *Burkholderia vietnamiensis G4* | bvi_Bcep1808_2152 | 240 |
| | | | | bvi_Bcep1808_2158 | 279 |
| | | | *Burkholderiales bacterium GJ-E10* | bbag_E1O_23150 | 169 |
| | | | *Caballeronia insecticola* | buo_BRPE64_ACDS04770 | 167 |
| | | | *Caldimonas brevitalea* | pbh_AAW51_0098 | 167 |
| | | | *Castellaniella defragrans* | cdn_BN940_08181 | 173 |
| | | | *Comamonas serinivorans* | cser_CCO03_18970 | 166 |
| | | | | cser_CCO03_18975 | 168 |
| | | | *Comamonas testosteroni TK102* | ctes_O987_14600 | 170 |
| | | | *Delftia lacustris* | dla_I6G47_20305 | 187 |
| | | | *Delftia sp. Cs1-4* | del_DelCs14_5662 | 166 |
| | | | *Delftia tsuruhatensis* | dts_BI380_20350 | 167 |
| | | | *Diaphorobacter ruginosibacter* | drg_H9K76_21515 | 177 |
| | | | *Diaphorobacter sp. HDW4A* | dih_G7047_25905 | 188 |
| | | | *Hydrogenophaga pseudoflava* | hpse_HPF_22800 | 172 |
| | | | | hpse_HPF_22805 | 175 |
| | | | *Hydrogenophaga sp. LPB0072* | hyl_LPB072_22445 | 180 |
| | | | | hyl_LPB072_22450 | 181 |
| | | | *Massilia oculi* | mtim_DIR46_18135 | 175 |
| | | | *Massilia sp. YMA4* | masy_DPH57_17525 | 166 |
| | | | *Mycetohabitans rhizoxinica* | brh_RBRH_01328 | 166 |
| | | | *Ottowia sp. oral taxon 894* | oto_ADJ79_06795 | 168 |
| | | | *Ottowia sp. oral taxon 894* | oto_ADJ79_06800 | 170 |
| | | | *Paenalcaligenes hominis* | phn_PAEH1_04210 | 168 |

| | | | | | |
|---|---|---|---|---|---|
| | | | *Paraburkholderia tropica* | ptro_G5S35_11920 | 167 |
| | | | *Pigmentiphaga aceris* | pacr_FXN63_19155 | 175 |
| | | | *Pigmentiphaga sp. H8* | pig_EGT29_07315 | 180 |
| | | | *Polaromonas naphthalenivorans* | pna_Pnap_4333 | 178 |
| | | | *Pulveribacter suum* | melm_C7H73_02965 | 181 |
| | | | *Pusillimonas sp. DMV24BSW_D* | pud_G9Q38_12345 | 171 |
| | | | *Pusillimonas sp. T7-7* | put_PT7_1417 | 176 |
| | | | *Rhodoferax sediminis Gr-4* | rhg_EXZ61_01095 | 168 |
| | | | *Simplicispira suum* | simp_C6571_08580 | 169 |
| | | | *Verminephrobacter eiseniae* | vei_Veis_4782 | 167 |
| | | Neisseriales | *Neisseria bacilliformis* | nbc_H3L91_00080 | 172 |
| | | | | nbc_H3L91_00085 | 173 |
| | | | | nbc_H3L91_00090 | 173 |
| | | | *Neisseria elongata* | nel_NELON_01840 | 167 |
| | | | | nel_NELON_01845 | 181 |
| | | | *Neisseria zalophi* | nzl_D0T92_10455 | 175 |
| | | | | nzl_D0T92_10460 | 167 |
| | | | *Simonsiella muelleri* | smur_BWP33_11975 | 166 |
| | | Nitrosomonadales | *Sideroxydans lithotrophicus* | slt_Slit_0538 | 172 |
| Firmicutes | 25 | Bacillales | *Anoxybacillus flavithermus* | afl_Aflv_0627 | 168 |
| | | | *Bacillus selenitireducens* | bse_Bsel_0234 | 191 |
| | | | *Geobacillus sp. LC300* | gel_IB49_04990 | 168 |
| | | | *Geobacillus stearothermophilus* | gse_GT50_04575 | 168 |
| | | | *Geobacillus subterraneus* | gsr_GS3922_03060 | 169 |
| | | Erysipelotrichales | *Intestinibaculum porci* | ebm_SG0102_16640 | 167 |
| | | Eubacteriales | *Acetivibrio saccincola* | hsc_HVS_10140 | 192 |
| | | | *Anaerostipes rhamnosivorans* | arf_AR1Y2_1992 | 280 |
| | | | *Clostridioides difficile 630* | cdf_CD630_35080 | 178 |
| | | | *Clostridium kluyveri DSM 555* | ckl_CKL_1112 | 166 |
| | | | *Clostridium perfringens 13* | cpe_CPE2284 | 170 |
| | | | *Eubacterium maltosivorans* | emt_CPZ25_010940 | 179 |
| | | | *Eubacterium siraeum 70/3* | esu_EUS_24460 | 210 |
| | | | | esu_EUS_24480 | 217 |
| | | | *Eubacterium siraeum V10Sc8a* | esr_ES1_06510 | 203 |
| | | | | esr_ES1_06520 | 210 |
| | | | *Geosporobacter ferrireducens* | gfe_Gferi_00415 | 234 |
| | | | *Hungateiclostridium thermocellum ATCC 27405* | cth_Cthe_1103 | 180 |
| | | | | cth_Cthe_1104 | 166 |
| | | | *Hungateiclostridium thermocellum DSM 1313* | ctx_Clo1313_1110 | 178 |
| | | | *Ruminococcus albus* | ral_Rumal_3060 | 237 |
| | | | *Ruminococcus bicirculans* | rus_RBI_I01475 | 169 |

| | | | | | |
|---|---|---|---|---|---|
| | | | *Ruminococcus champanellensis* | rch_RUM_03620 | 195 |
| | | | | rch_RUM_16890 | 173 |
| | | | *Thermoclostridium stercorarium DSM 8532* | css_Cst_c12640 | 173 |
| Cyanobacteria | 29 | Chroococcales | *Crocosphaera subtropica* | cyt_cce_2840 | 180 |
| | | | *Gloeothece citriformis* | cyc_PCC7424_0260 | 167 |
| | | Gloeobacterales | *Gloeobacter kilaueensis* | glj_GKIL_0446 | 185 |
| | | | | glj_GKIL_4335 | 172 |
| | | | *Gloeobacter violaceus* | gvi_gll2255 | 171 |
| | | | | gvi_glr3426 | 186 |
| | | Nostocales | *Calothrix sp. PCC 6303* | calt_Cal6303_362 | 183 |
| | | | *Cylindrospermum stagnale* | csg_Cylst_2769 | 174 |
| | | | *Nostoc flagelliforme* | nfl_COO91_06898 | 168 |
| | | Oscillatoriales | *Geitlerinema sp. PCC 7407* | gei_GEI7407_3664 | 169 |
| | | | *Microcoleus sp. PCC 7113* | mic_Mic7113_5040 | 174 |
| | | | *Moorea producens* | mpro_BJP34_19960 | 175 |
| | | Pseudoanabaenales | *Leptolyngbya sp. NIES-3755* | len_LEP3755_39030 | 176 |
| | | | *Leptolyngbya sp. PCC 7376* | lep_Lepto7376 | 243 |
| | | | *Pseudanabaena sp. ABRG5-3* | pser_ABRG53_0283 | 172 |
| | | | *Pseudanabaena sp. PCC 7367* | pseu_Pse7367_2053 | 177 |
| | | | | pseu_Pse7367_2054 | 166 |
| | | | *Halomicronema hongdechloris* | hhg_XM38_026330 | 173 |
| | | | *Thermosynechococcus elongatus* | tel_tlr0682_ 172 | 172 |
| | | | *Thermosynechococcus sp. NK55* | thn_NK55_01305 | 172 |
| | | Synechococcales | *Chamaesiphon minutus* | cmp_Cha6605_2755 | 168 |
| | | | *Cyanobium gracile* | cgc_Cyagr_1414 | 169 |
| | | | *Prochlorococcus marinus MIT 9211* | pmj_P9211_15291 | 171 |
| | | | *Prochlorococcus marinus MIT 9301* | pmg_P9301_07011 | 175 |
| | | | *Prochlorococcus marinus MIT 9312* | pmi_PMT9312_1201 | 177 |
| | | | *Synechococcus sp. PCC 7003* | syl_AWQ21_14105 | 173 |
| | | | *Synechococcus sp. PCC 73109* | syv_AWQ23_13975 | 212 |
| | | | *Synechococcus sp. PCC 7502* | synp_Syn7502_00935 | 198 |
| | | | *Synechococcus sp. PCC 7002* | syp_SYNPCC7002_A2804 | 170 |
| Actinobacteria | 4 | Bifidobacteriales | *Bifidobacterium thermophilum* | btp_D805_0701 | 187 |
| | | Corynebacteriales | *Corynebacterium callunae* | ccn_H924_06030 | 224 |
| | | Eggerthellales | *Gordonibacter pamelaeae* | gpa_GPA_11520 | 240 |
| | | Euzebyales | *Euzebya pacifica DY32-46* | euz_DVS28_b0477 | 227 |
| Deinococcus-Thermus | 1 | Thermales | *Thermus sp. CCB_US3_UF1* | thc_TCCBUS3UF1_3990 | 176 |
| Thermodesulfobacteria | 7 | Desulfobulbales | *Desulfobulbus oralis* | deo_CAY53_11360 | 200 |
| | | | *Desulfurivibrio alkaliphilus* | dak_DaAHT2_2283 | 182 |
| | | Desulfuromonadales | *Desulfuromonas soudanensis* | des_DSOUD_2157 | 193 |

| | | | | | |
|---|---|---|---|---|---|
| | | | *Pelobacter carbinolicus* | pca_Pcar_2143 | 170 |
| | | Geobacterales | *Geobacter daltonii FRC-32* | geo_Geob_3369 | 208 |
| | | | *Geobacter uraniireducens* | gur_Gura_2677 | 193 |
| | | Desulfomonilales | *Desulfomonile tiedjei* | dti_Desti_4024 | 204 |
| Myxococcota | 39 | Myxococcales | *Anaeromyxobacter dehalogenans 2CP-1* | acp_A2cp1_0669 | 188 |
| | | | *Anaeromyxobacter sp. Fw109-5* | afw_Anae109_0680 | 191 |
| | | | *Archangium gephyra* | age_AA314_07645 | 202 |
| | | | | age_AA314_08283 | 204 |
| | | | *Chondromyces crocatus* | ccro_CMC5_052100 | 182 |
| | | | | ccro_CMC5_052110 | 188 |
| | | | | ccro_CMC5_052120 | 193 |
| | | | *Corallococcus coralloides* | ccx_COCOR_06280 | 198 |
| | | | *Corallococcus macrosporus* | mfu_LILAB_36405 | 203 |
| | | | *Cystobacter fuscus* | cfus_CYFUS_00131 | 202 |
| | | | | cfus_CYFUS_00190 | 202 |
| | | | | cfus_CYFUS_00751 | 201 |
| | | | *Haliangium ochraceum* | hoh_Hoch_4154 | 175 |
| | | | | hoh_Hoch_4155 | 177 |
| | | | *Labilithrix luteola* | llu_AKJ09_08823 | 185 |
| | | | | llu_AKJ09_08824 | 166 |
| | | | *Melittangium boletus* | mbd_MEBOL_001002 | 201 |
| | | | | mbd_MEBOL_001435 | 203 |
| | | | *Minicystis rosea* | mrm_A7982_03796 | 187 |
| | | | | mrm_A7982_03797 | 173 |
| | | | *Myxococcus fulvus* | mfb_MFUL124B02 | 183 |
| | | | *Myxococcus hansupus* | mym_A176_001126 | 217 |
| | | | *Myxococcus macrosporus* | mmas_MYMAC_00557 | 205 |
| | | | *Myxococcus stipitatus* | msd_MYSTI_06366 | 184 |
| | | | *Myxococcus xanthus* | mxa_MXAN_5783 | 208 |
| | | | *Sandaracinus amylolyticus* | samy_DB32_004027 | 176 |
| | | | | samy_DB32_004028 | 176 |
| | | | *Sorangium cellulosum So ce56* | scl_sce4273 | 179 |
| | | | | scl_sce4274 | 192 |
| | | | | scl_sce4275 | 192 |
| | | | *Sorangium cellulosum So0157-2* | scu_SCE1572_2584 | 178 |
| | | | | scu_SCE1572_2585 | 193 |
| | | | *Stigmatella aurantiaca* | sur_STAUR_0003 | 202 |
| | | | | sur_STAUR_0004 | 203 |
| | | | | sur_STAUR_1125 | 203 |
| | | | | sur_STAUR_6449 | 205 |
| | | | | sur_STAUR_6450 | 204 |
| | | | | sur_STAUR_6924 | 204 |

| | | | | | |
|---|---|---|---|---|---|
| | | | *Vulgatibacter incomptus* | vin_AKJ08_0698 | 171 |
| Verrucomicrobiota | 4 | Opitutales | *Nibricoccus aquaticus HZ-65* | vbh_CMV30_12470 | 166 |
| | | | | vbh_CMV30_16935 | 210 |
| | | Verrucomicrobiales | *Luteolibacter luteus* | luo_HHL09_15680 | 188 |
| | | | *Roseimicrobium sp. ORNL1* | roo_G5S37_24550 | 223 |
| Aquificae | 1 | Desulfurobacteriales | *Thermovibrio ammonificans* | tam_Theam_1352 | 185 |
| Acidobacteriota | 1 | Acidobacteriales | *Koribacter versatilis* | aba_Acid345_2448 | 180 |
| Proteobacteria-Acidithiobacilli | 1 | Acidithiobacillaes | *Acidithiobacillus ferrivorans* | afi_Acife_1976 | 190 |
| Bdellovibrionota | 11 | Bacteriovoracales | *Bacteriovorax stolpii* | bsto_C0V70_00655 | 170 |
| | | | *Halobacteriovorax marinus* | bmx_BMS_0122 | 183 |
| | | | *Halobacteriovorax sp. BALOs_7* | hax_BALOs_0146 | 176 |
| | | Bdellovibrionales | *Bdellovibrio bacteriovorus 109J* | bbac_EP01_16385 | 180 |
| | | | *Bdellovibrio bacteriovorus HD100* | bba_Bd1290 | 179 |
| | | | *Bdellovibrio bacteriovorus Tiberius* | bbat_Bdt_1269 | 186 |
| | | | *Bdellovibrio bacteriovorus W* | bbw_BDW_04435 | 189 |
| | | | *Bdellovibrio exovorus* | bex_A11Q_1638 | 181 |
| | | | *Bdellovibrio sp. NC01* | bdc_DOE51_05870 | 186 |
| | | | *Bdellovibrio sp. qaytius* | bdq_CIK05_05425 | 204 |
| | | | *Bdellovibrio sp. ZAP7* | bdz_DOM22_13925 | 177 |
| Proteobacteria-Alpha | 2 | no rank | *Alpha proteobacterium HIMB5* | apm_HIMB5_000081 | 195 |
| | | | *Puniceispirillum marinum (Candidatus)* | apb_SAR116_2527 | 188 |
| Nitrospirota | 3 | Nitrospirales | *Nitrospira defluvii* | nde_NIDE3438 | 169 |
| | | | | nde_NIDE3439 | 166 |
| | | | *Nitrospira inopinata (Candidatus)* | nio_NITINOP_2648 | 196 |
| Armatimonadota | 1 | Chthonomonadales | *Chthonomonas calidirosea* | ccz_CCALI_00192 | 176 |

Detailed information of the 226 large major pilins in regard of corresponding phylum or proteobacterial class (1st row), number of total sequences in that subset (2nd row), bacterial order (3rd row), bacterial species (4th row), locus tag (fifth row) and major pilin length in aa (last row).

**Table S3. *M. xanthus* and *E. coli* strains used in this work.**

| Strain | Description/Genotype[1] | Reference or source |
|---|---|---|
| ***M. xanthus*** | | |
| DK1622 | wildtype | [8] |
| DK10410 | Δ*pilA* (ΔMXAN_5783) | [9] |
| DK10409 | Δ*pilT* (ΔMXAN_5787) | [9] |
| SA11429 | *pilA*::*pilA*-R30Q (pMAT473) | This study |
| SA11431 | *pilA*::*pilA*-R30A (pMAT474) | This study |
| SA11435 | *pilA*::*pilA*-K37Q (pMAT476) | This study |
| SA11437 | *pilA*::*pilA*-K37A (pMAT477) | This study |
| SA11439 | *pilA*::*pilA*- E53Q (pMAT479) | This study |
| SA11441 | *pilA*::*pilA*- E53A (pMAT480) | This study |
| SA11451 | *pilA*::*pilA*-D55N (pMAT485) | This study |
| SA11453 | *pilA*::*pilA*-D55A (pMAT486) | This study |
| SA11457 | *pilA*::*pilA*-R73Q (pMAT488) | This study |
| SA11459 | *pilA*::*pilA*-R73A (pMAT489) | This study |
| SA11463 | *pilA*::*pilA*-R109Q (pMAT491) | This study |
| SA11491 | *pilA*::*pilA*-R109A (pMAT492) | This study |
| SA11469 | *pilA*::*pilA*-K48Q (pMAT494) | This study |
| SA11471 | *pilA*::*pilA*-K48A (pMAT495) | This study |
| SA11475 | *pilA*::*pilA*-E69Q (pMAT497) | This study |
| SA11477 | *pilA*::*pilA*-E69A (pMAT498) | This study |
| SA11497 | *pilA*::*pilA*-R70Q (pMAT528) | This study |
| SA11499 | *pilA*::*pilA*-R70A (pMAT505) | This study |
| SA11430 | *pilA*::*pilA*-R30Q (pMAT473) | This study |
| SA11432 | Δ*pilT*; *pilA*::*pilA*-R30A (pMAT474) | This study |
| SA11436 | Δ*pilT*; *pilA*::*pilA*-K37Q (pMAT476) | This study |

| | | |
|---|---|---|
| SA11485 | Δ*pilT*; *pilA*::*pilA*-K37A (pMAT477) | This study |
| SA11440 | Δ*pilT*; *pilA*::*pilA*- E53Q (pMAT479) | This study |
| SA11442 | Δ*pilT*; *pilA*::*pilA*- E53A (pMAT480) | This study |
| SA11452 | Δ*pilT*; *pilA*::*pilA*-D55N (pMAT485) | This study |
| SA11454 | Δ*pilT*; *pilA*::*pilA*-D55A (pMAT486) | This study |
| SA11458 | Δ*pilT*; *pilA*::*pilA*-R73Q (pMAT488) | This study |
| SA11460 | Δ*pilT*; *pilA*::*pilA*-R73A (pMAT489) | This study |
| SA11464 | Δ*pilT*; *pilA*::*pilA*-R109Q (pMAT491) | This study |
| SA11466 | Δ*pilT*; *pilA*::*pilA*-R109A (pMAT492) | This study |
| SA11470 | Δ*pilT*; *pilA*::*pilA*-K48Q (pMAT494) | This study |
| SA11472 | Δ*pilT*; *pilA*::*pilA*-K48A (pMAT495) | This study |
| SA11476 | Δ*pilT*; *pilA*::*pilA*-E69Q (pMAT497) | This study |
| SA11478 | Δ*pilT*; *pilA*::*pilA*-E69A (pMAT498) | This study |
| SA12322 | Δ*pilT*; *pilA*::*pilA*-R70Q (pMAT528) | This study |
| SA12300 | Δ*pilT*; *pilA*::*pilA*-R70A (pMAT505) | This study |
| ***E. coli*** | | |
| NEB® Turbo | F' *proA*$^{+}$*B*$^{+}$ *lacI*$^{q}$ Δ*lacZM15 / fhuA2* Δ*(lac-proAB) glnV galK16 galE15 R(zgb-210::Tn10)*Tet$^{S}$ *endA1 thi-1* Δ*(hsdS-mcrB)5* | New England Biolabs |

[1] Plasmids used for construction of *pilA*-variants at the endogenous locus are indicated in brackets.

**Table S4. Plasmids used in this work.**

| **Plasmids** | | |
|---|---|---|
| pBJ114 | *galK* containing vector for generation of in-frame deletions in *M. xanthus*, Kan$^R$ | [10] |
| pMAT436 | pBJ114, contains 2594 bp gene fragment of *pilA* region with *pilA* gene from 941-1603 bp | This study |
| pMAT473 | pMAT436, construct for endogenous *pilA*-R30Q mutation | This study |
| pMAT474 | pMAT436, construct for endogenous *pilA*-R30A mutation | This study |
| pMAT476 | pMAT436, construct for endogenous *pilA*-K37Q mutation | This study |
| pMAT477 | pMAT436, construct for endogenous *pilA*-K37A mutation | This study |
| pMAT479 | pMAT436, construct for endogenous *pilA*- E53Q mutation | This study |
| pMAT480 | pMAT436, construct for endogenous *pilA*- E53A mutation | This study |
| pMAT485 | pMAT436, construct for endogenous *pilA*-D55N mutation | This study |
| pMAT486 | pMAT436, construct for endogenous *pilA*-D55A mutation | This study |
| pMAT488 | pMAT436, construct for endogenous *pilA*-R73Q mutation | This study |
| pMAT489 | pMAT436, construct for endogenous *pilA*-R73A mutation | This study |
| pMAT491 | pMAT436, construct for endogenous *pilA*-R109Q mutation | This study |
| pMAT492 | pMAT436, construct for endogenous *pilA*-R109A mutation | This study |
| pMAT494 | pMAT436, construct for endogenous *pilA*-K48Q mutation | This study |
| pMAT495 | pMAT436, construct for endogenous *pilA*-K48A mutation | This study |
| pMAT497 | pMAT436, construct for endogenous *pilA*-E69Q mutation | This study |
| pMAT498 | pMAT436, construct for endogenous *pilA*-E69A mutation | This study |
| pMAT528 | pMAT436, construct for endogenous *pilA*-R70Q mutation | This study |
| pMAT505 | pMAT436, construct for endogenous *pilA*-R70A mutation | This study |

**Table S5. Oligonucleotides used in this work.**

| Name | Sequence[1] |
|---|---|
| *pilA*-up EcoRI+ | **GCGC<u>GAATTC</u>**CACTGGCGCGACCACCGAC |
| *pilA*-down HindIII- | **GCGC<u>AAGCTT</u>**GAACTGAATGCCACCCGCC |
| *pilA*-E2 | TTGAACGAGGGGACGCTGGA |
| *pilA*-F3 | TGGCCTGGGGCAATCTCAAG |
| *pilA*-G | CCTGGCCGCCATCGCCATCC |
| *pilA*-H | CGATCACCCAGTCATCGAAG |
| *pilA*-stop- | TTACTGGGCCGCGCCGTC |
| *pilA*-start+ | ATGCGCGTCTCGCGATTC |
| M13 rev | GAGCGGATAACAATTTCACACAGG |
| M13 forw | AGGGTTTTCCCAGTCACGACGTT |
| pilA-R42(30)Q+ | TTCATCAAGTTCCAGGCCCAGTCGAAGCAGTCCGAGGCG |
| pilA-R42(30)Q- | CGCCTCGGACTGCTTCGACTGGGCCTGGAACTTGATGAA |
| pilA-R42(30)A+ | TTCATCAAGTTCCAGGCCGCCTCGAAGCAGTCCGAGGCG |
| pilA-R42(30)A- | CGCCTCGGACTGCTTCGAGGCGGCCTGGAACTTGATGAA |
| pilA-K49(37)Q+ | TCGAAGCAGTCCGAGGCGCAGACGAACCTCAAGGCGCTG |
| pilA-K49(37)Q- | CAGCGCCTTGAGGTTCGTCTGCGCCTCGGACTGCTTCGA |
| pilA-K49(37)A+ | TCGAAGCAGTCCGAGGCGGCCACGAACCTCAAGGCGCTG |
| pilA-K49(37)A- | CAGCGCCTTGAGGTTCGTGGCCGCCTCGGACTGCTTCGA |
| pilA-E65(53)Q+ | CAGAAGTCGTTCTTCTCCCAGAAGGACCGTTACTCCGAC |
| pilA-E65(53)Q- | GTCGGAGTAACGGTCCTTCTGGGAGAAGAACGACTTCTG |
| pilA-E65(53)A+ | CAGAAGTCGTTCTTCTCCGCCAAGGACCGTTACTCCGAC |
| pilA-E65(53)A- | GTCGGAGTAACGGTCCTTGGCGGAGAAGAACGACTTCTG |
| pilA-D67(55)N+ | TCGTTCTTCTCCGAGAAGAACCGTTACTCCGACTTCGCC |
| pilA-D67(55)N- | GGCGAAGTCGGAGTAACGGTTCTTCTCGGAGAAGAACGA |
| pilA-D67(55)A+ | TCGTTCTTCTCCGAGAAGGCGCGTTACTCCGACTTCGCC |

| | |
|---|---|
| pilA-D67(55)A- | GGCGAAGTCGGAGTAACGCGCCTTCTCGGAGAAGAACGA |
| pilA-R85(73)Q+ | GCGCCGGAGCGCGGCAACCAGTACGGCTACCGTGTGTCC |
| pilA-R85(73)Q- | GGACACACGGTAGCCGTACTGGTTGCCGCGCTCCGGCGC |
| pilA-R85(73)A+ | GCGCCGGAGCGCGGCAACGCGTACGGCTACCGTGTGTCC |
| pilA-R85(73)A- | GGACACACGGTAGCCGTACGCGTTGCCGCGCTCCGGCGC |
| pilA R121(109)Q+ | ATCTCCAACGACTCGTTCCAGTTCGGTGCCAACAGCGCC |
| pilA R121(109)Q- | GGCGCTGTTGGCACCGAACTGGAACGAGTCGTTGGAGAT |
| pilA R121(109)A+ | ATCTCCAACGACTCGTTCGCGTTCGGTGCCAACAGCGCC |
| pilA R121(109)A- | GGCGCTGTTGGCACCGAACGCGAACGAGTCGTTGGAGAT |
| pilA K60(48)Q+ | GCGCTGTACACCGCGCAGCAGTCGTTCTTCTCCGAGAAG |
| pilA K60(48)Q- | CTTCTCGGAGAAGAACGACTGCTGCGCGGTGTACAGCGC |
| pilA K60(48)A+ | GCGCTGTACACCGCGCAGGCGTCGTTCTTCTCCGAGAAG |
| pilA K60(48)A- | CTTCTCGGAGAAGAACGACGCCTGCGCGGTGTACAGCGC |
| pilA-E81(69)Q+ | GAAATCGGCTTCGCGCCGCAGCGCGGCAACCGTTACGGC |
| pilA-E81(69)Q- | GCCGTAACGGTTGCCGCGCTGCGGCGCGAAGCCGATTTC |
| pilA-E81(69)A+ | GAAATCGGCTTCGCGCCGGCGCGCGGCAACCGTTACGGC |
| pilA-E81(69)A- | GCCGTAACGGTTGCCGCGCGCCGGCGCGAAGCCGATTTC |
| pilA-R82(70)Q+ | ATCGGCTTCGCGCCGGAGCAGGGCAACCGTTACGGCTAC |
| pilA-R82(70)Q- | GTAGCCGTAACGGTTGCCCTGCTCCGGCGCGAAGCCGAT |
| pilA-R82(70)A+ | ATCGGCTTCGCGCCGGAGGCGGGCAACCGTTACGGCTAC |
| pilA-R82(70)A- | GTAGCCGTAACGGTTGCCCGCCTCCGGCGCGAAGCCGAT |

[1] Sequences added for cloning purposes are indicated in bold. Restriction sites are underlined.

### Supplementary References


1 Treuner-Lange, A. *et al.* PilY1 and minor pilins form a complex priming the type IVa pilus in *Myxococcus xanthus*. *Nat Commun* **11**, 5054, (2020).

2 Kolappan, S. *et al.* Structure of the *Neisseria meningitidis* type IV pilus. *Nat Commun* **7**, 13015, (2016).

3 Wang, F. *et al.* Cryoelectron microscopy reconstructions of the *Pseudomonas aeruginosa* and *Neisseria gonorrhoeae* type IV pili at sub-nanometer resolution. *Structure* **25**, 1423-1435 e1424, (2017).

4 Neuhaus, A. *et al.* Cryo-electron microscopy reveals two distinct type IV pili assembled by the same bacterium. *Nat Commun* **11**, 2231, (2020).

5 Gu, Y. *et al.* Structure of *Geobacter* pili reveals secretory rather than nanowire behaviour. *Nature* **597**, 430-434, (2021).

6 Bardiaux, B. *et al.* Structure and assembly of the enterohemorrhagic *Escherichia coli* type 4 pilus. *Structure* **27**, 1082-1093 e1085, (2019).

7 Lopez-Castilla, A. *et al.* Structure of the calcium-dependent type 2 secretion pseudopilus. *Nat Microbiol* **2**, 1686-1695, (2017).

8 Kaiser, D. Social gliding is correlated with the presence of pili in *Myxococcus xanthus*. *Proc. Natl. Acad. Sci. USA* **76**, 5952-5956, (1979).

9 Wu, S. S., Wu, J. & Kaiser, D. The *Myxococcus xanthus pilT* locus is required for social gliding motility although pili are still produced. *Mol. Microbiol.* **23**, 109-121, (1997).

10 Julien, B., Kaiser, A. D. & Garza, A. Spatial control of cell differentiation in *Myxococcus xanthus*. *Proc. Natl. Acad. Sci. USA* **97**, 9098-9103, (2000).
